# Pathways to polar adaptation in fishes revealed by long-read sequencing

**DOI:** 10.1101/2021.11.12.468413

**Authors:** Scott Hotaling, Thomas Desvignes, John S. Sproul, Luana S.F. Lins, Joanna L. Kelley

## Abstract

Long-read sequencing is driving a new reality for genome science where highly contiguous assemblies can be produced efficiently with modest resources. Genome assemblies from long-read sequences are particularly exciting for understanding the evolution of complex genomic regions that are often difficult to assemble. In this study, we leveraged long-read sequencing data to generate a high-quality genome assembly for an Antarctic eelpout, *Opthalmolycus amberensis*, the first for the globally distributed family Zoarcidae. We used this assembly to understand how *O. amberensis* has adapted to the harsh Southern Ocean and compared it to another group of Antarctic fishes: the notothenioids. We showed that selection has largely acted on different targets in eelpouts relative to notothenioids. However, we did find some overlap; in both groups, genes involved in membrane structure, thermal tolerance, and vision have evidence of selection. We found evidence for historical shifts of transposable element activity in *O. amberensis* and other polar fishes, perhaps reflecting a response to environmental change. We were specifically interested in the evolution of two complex genomic loci known to underlie key adaptations to polar seas: hemoglobin and antifreeze proteins (AFPs). We observed unique evolution of the hemoglobin MN cluster in eelpouts and related fishes in the suborder Zoarcoidei relative to other Perciformes. For AFPs, we identified the first species in the suborder with no evidence of *afpIII* sequences (*Cebidichthys violaceus*) in the genomic region where they are found in all other Zoarcoidei, potentially reflecting a lineage-specific loss of this cluster. Beyond polar fishes, our results highlight the power of long-read sequencing to understand genome evolution.

## Introduction

Long-reads are revolutionizing genome sequencing by allowing for improved resolution of genotypes across the tree of life (Chawla et al., 2021; Hoencamp et al., 2021; Hotaling, Kelley, & Frandsen, 2021; Hotaling, Sproul, et al., 2021; Sone et al., 2019). This new potential is particularly valuable for understanding phenotypes that stem from complex, difficult to assemble regions of the genome (e.g., highly repetitive regions). For instance, the evolution of antifreeze proteins (AFPs)—proteins that adsorb to ice and limit its growth (Davies, Baardsnes, Kuiper, & Walker, 2002; X. Li et al., 1985)—has led to a wide array of teleost fishes thriving in marine habitats that are commonly below freezing and ice-laden (Hobbs, Hall, Graham, Davies, & Fletcher, 2020). AFPs are commonly found in highly repetitive, tandem arrays in the genome and have experienced a large amount of duplication, rearrangement, and modification (e.g., Deng, Cheng, Ye, He, & Chen, 2010; Scott, Hew, & Davies, 1985). Thus, the evolution of AFPs and phenotypes with similarly complex genomic underpinnings has been difficult to understand despite their importance for many ecological communities.

Extreme environments provide insight into the nature of adaptive evolution due to their strong, and often long-term, selective pressures (Kim et al., 2019; S. Xu, Wang, Guo, He, & Shi, 2020). Extreme habitats are even more useful for evolutionary inquiry when distantly related lineages are subjected to the same selective pressures and their responses are compared. Under this framework, it is possible to disentangle lineage-specific evolutionary responses from more general patterns (e.g., convergent evolution). The Southern Ocean surrounding Antarctica is one of the most contiguous extreme environments on Earth. It is generally recognized as latitudes south of the Antarctic Convergence (also known as Antarctic Polar Front), where extremely cold (sea surface temperature < 0°C), northward-flowing water from Antarctica meets the comparatively warmer waters of the subantarctic (sea surface temperature > 5.5°C; Moore, Abbott, & Richman, 1999). The extreme nature of the Southern Ocean largely stems from three factors: chronic subfreezing temperatures, high levels of dissolved oxygen, and dramatic seasonal shifts in light availability (DeVries & Steffensen, 2005). Such extreme conditions greatly influence the biological diversity of the Southern Ocean (Griffiths, 2010) and have driven the evolution of an array of extreme phenotypes (e.g., sea spider gigantism, A. L. Moran & Woods, 2012).

Among fishes, the most well-known outcome of the long-term selection at both poles is the evolution of AFPs. Based on protein structure, multiple different AFP types have evolved in fishes, including antifreeze glycoproteins (AFGPs) in cod and Antarctic notothenioids and Type III antifreeze proteins in eelpouts (Davies et al., 2002; X. Li et al., 1985). AFPs are common in polar fishes but knowledge of their molecular evolution remains incomplete (but see Chen, DeVries, & Cheng, 1997a; Chen, DeVries, & Cheng, 1997b; Deng et al., 2010; Zhuang, Yang, Murphy, & Cheng, 2019). Similarly, hemoglobin evolution appears to be an interesting component of thermal adaptation for fishes, including among the white-blooded Antarctic icefishes (family Channichthyidae, Ruud, 1954), which are the only vertebrates known to live without hemoglobin (Beck et al., 2021; Kim et al., 2019). To date, practical limitations have inhibited our understanding of AFP and hemoglobin evolution, particularly the challenge of assembling these repeat-rich, complex loci with short-read sequence data which cannot span repetitive regions and therefore lack the power to reveal higher-level genomic architecture (e.g., Ahn et al., 2017).

Thus far, genomic investigation of Antarctic fishes has been restricted to the notothenioid adaptive radiation (e.g., Kim et al., 2019; Near et al., 2012). These studies have increased understanding of how cryonotothenioids (Antarctic notothenioids) have adapted to the Southern Ocean (Chen et al., 2019) and how the unique white-blooded icefishes evolved following the loss of hemoglobin (Near, Parker, & Detrich III, 2006). Still, many questions remain. Namely, how generalizable is the evolutionary trajectory of cryonotothenioids relative to other fishes in the Southern Ocean? With roughly ∼300 described species distributed around the world and many in Antarctica (Anderson, 1994; Hotaling, Borowiec, Lins, Desvignes, & Kelley, 2021; Møller, Nielsen, & Anderson, 2005), the eelpout family (family Zoarcidae) is the most speciose family within the suborder Zoarcoidei (also referred to as the infraorder Zoarcales). Similar to cryonotothenioids, eelpouts have evolved AFPs to mitigate freezing stress, specifically Type III AFPs (*afpIII*; X. Li et al., 1985). The most recent common ancestor of eelpouts and cryonotothenioids diverged ∼80 million years ago (Rabosky et al., 2018)—long before the cooling of the Southern Ocean—making these groups an ideal system for studying potential convergent evolution to the contemporary, subfreezing conditions of polar seas.

In this study, we used long-read sequencing to generate a high-quality genome assembly for an Antarctic eelpout, *Ophthalmolycus amberensis*, the first for the family Zoarcidae. We then leveraged our new genomic resource to better understand how fishes have adapted to the Southern Ocean through whole-genome perspectives on genome evolution and fine-scale investigation of synteny and copy number variation in key adaptive regions (i.e., hemoglobin and AFPs). Collectively, our results highlight convergent and species-specific mechanisms underlying the adaptation of fishes to the harsh conditions of the Southern Ocean. Our study also provides practical evidence for the power and promise of long-read sequencing for understanding genome biology, particularly for highly repetitive and/or complex genomic regions.

## Materials and methods

### Species description and sample collection

*Ophthalmolycus amberensis* is a coastal Antarctic species, likely with a circumpolar distribution (Fig. 1). *Ophthalmolycus amberensis* individuals were collected from a depth of 360-380 meters in May 2016, using a 5-feet Blake trawl deployed from the Antarctic Research and Supply Vessel (ARSV) *Laurence M. Gould* in the Gerlache Strait (64°44’S, 63°01’W) on the West Antarctic Peninsula. Live fish were transferred immediately from the trawl net to the ship aquaria for transport to the aquatic facilities at Palmer Station, Antarctica, where they were maintained in flow-through seawater tanks at ∼0°C. Following euthanasia with a lethal dose of MS-222 (Syndel, Ferndale, WA, USA), individuals were dissected and sampled. Skeletal muscle was flash frozen in liquid nitrogen and stored at -80°C, fin clips were stored in 80% ethanol at room temperature, and liver was preserved in RNALater at 4°C until it was moved to longer term - 80°C storage. All procedures were performed according to protocols approved by the Institutional Animal Care and Use Committees (IACUC) of the University of Oregon (#13-27RRAA). We confirmed visual species identification through sequencing of the *mitochondrial cytochrome c oxidase I* (*mt-co1*) locus and comparison to sequences on GenBank.

**Figure 1.**
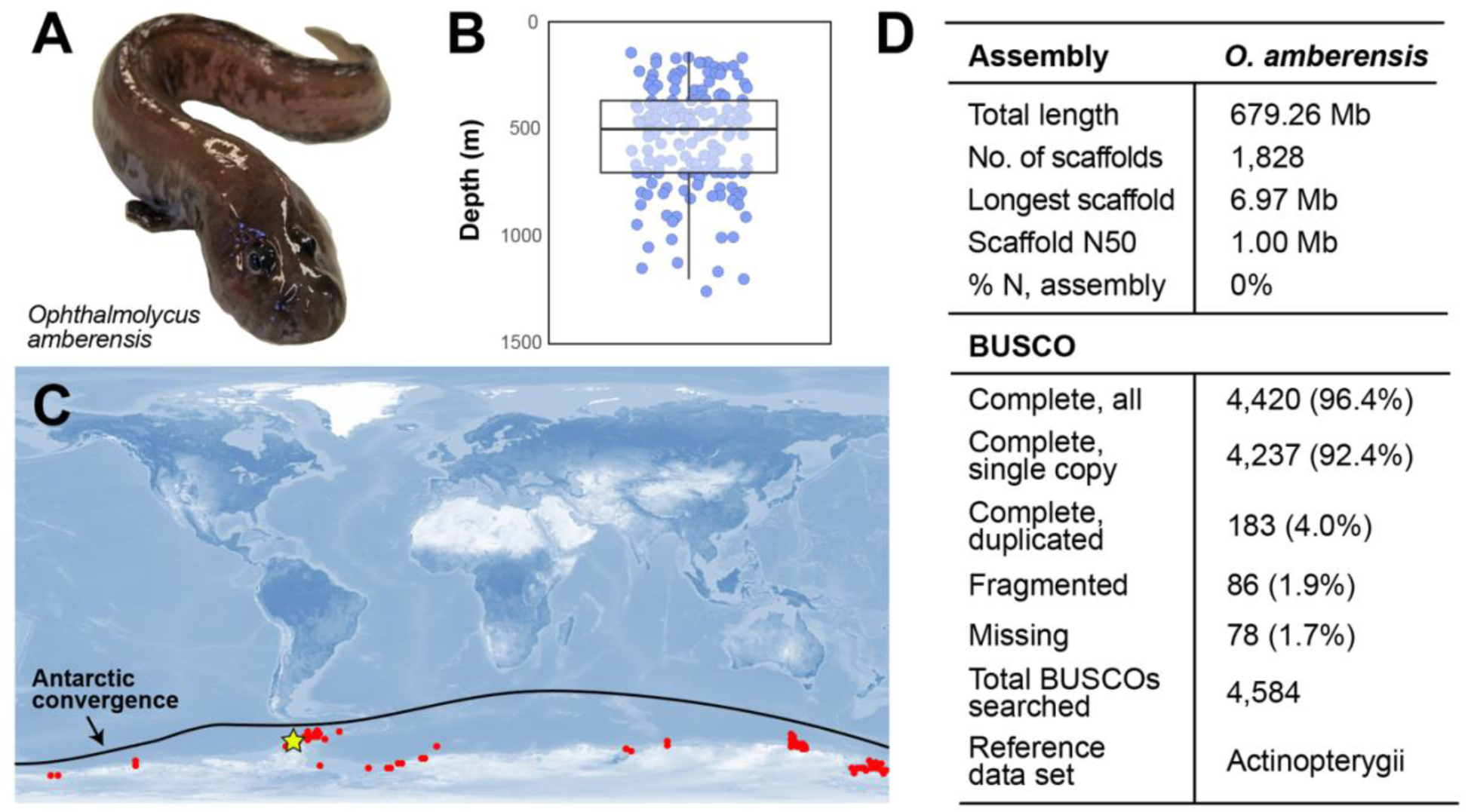
(A) An Antarctic eelpout, *Ophthalmolycus amberensis*, sampled from the Gerlache Strait on the West Antarctic Peninsula. (B) Depth records (*n* = 168 records) for *O. amberensis* from GBIF (19 October 2021, GBIF occurrence download: https://doi.org/10.15468/dl.uu7w49). (C) The distribution of *O. amberensis* from AquaMaps (https://www.aquamaps.org/). Red circles represent sampling localities and the dark line indicates the Antarctic Convergence (or Antarctic Polar Front). A yellow star denotes where the specimens used in this study were collected. (D) Assembly and BUSCO (Benchmarking Universal Single-Copy Orthologs; Simão, Waterhouse, Ioannidis, Kriventseva, & Zdobnov, 2015) statistics for the *O. amberensis* genome assembly.

### Sequence data collection

We collected three sequence data sets for this study: long-read Pacific Bioscience (PacBio), short-read Illumina whole-genome (DNA-seq), and short-read Illumina transcriptomic (RNA-seq). Our sequence data were collected from three individuals and tissue types: PacBio (EP06, muscle), DNA-seq (EP10, fin clip), and RNA-seq (EP01, liver). PacBio Sequel SMRTbell libraries were prepared and sequenced at the Oregon State University Center for Quantitative Life Sciences on a PacBio Sequel platform with eight SMRT cells. For DNA-seq library preparation, genomic DNA was extracted using a MagAttract HMW DNA kit and the library was prepared following an Illumina PCR-free protocol with a 400-bp insert size. The resulting DNA-seq library was sequenced at MedGenome, Inc. on an Illumina HiSeq 4000 platform with 150-bp paired-end sequencing chemistry. Total RNA was extracted from ∼30 mg of liver tissue using the Macherey-Nagel NucleoSpin® kit and the manufacturer protocol. The tissue was flash frozen with liquid nitrogen and crushed to homogenize it before extraction. An RNA-seq library was prepared by the University of Utah Genomics Core Facility and sequenced on an Illumina HiSeq 2500 with 125-bp paired-end sequencing chemistry. We assessed sequence quality for the DNA-seq and RNA-seq data sets with FastQC v0.11.4 (Andrews, 2010). We trimmed the raw DNA-seq and RNA-seq reads with Trim Galore! v0.4.2 (Krueger, 2015) with stringency of 5, quality of 28, and a minimum length of 50 bp.

### Genome assembly

We first estimated the genome size and heterozygosity of the *O. amberensis* genome for the DNA-seq data using the online portal of GenomeScope (Vurture et al., 2017). We produced the input for GenomeScope, a histogram of 21-mer counts, with jellyfish v2.2.5 (Melsted & Pritchard, 2011). For genome assembly, we first combined our separate subread BAM files into a combined BAM using ‘pbmerge’, from the PacBio toolkit. Next, using ‘bam2fastx’, another PacBio tool, we converted the raw, combined subread BAM into FASTQ format for downstream analyses. Scripts and commands used in this study are provided on GitHub (https://github.com/scotthotaling/oamberensis_genome).

We assembled the genome from PacBio sequences only using Canu v1.8 (Koren et al., 2017), a dedicated long-read assembler designed for PacBio data. We changed one parameter, cnsErrorRate = 0.25, all other parameters were left as defaults. We also specified an expected genome size of 700 Mb. For all iterations of the genome assembly, we generated summary statistics using the Assemblathon2 summary script (assemblathon_stats.pl, Bradnam et al., 2013). We also assessed completeness by calculating the number of observed single-copy orthologs (BUSCOs) in the assembly using BUSCO v3 and the 4,584 “actinopterygii_odb9” reference gene set (Simão et al., 2015).

The initial Canu assembly (v1) of the *O. amberensis* genome was ∼40% larger than expected at 1.02 Gb with many duplicated regions (*n* = 1180 complete and duplicated BUSCOs), likely due to heterozygosity between haplotypes. To resolve duplicate contigs we used Purge Haplotigs (Roach et al. 2018). We first mapped raw PacBio reads to the v1 assembly with Minimap2 v2.14-r883 (Li 2018) and the options: -ax -map-pb. Next, we sorted the resulting BAM file of aligned reads with samtools v1.9 (Li et al. 2009) and default settings. We used the ‘readhist’ function of Purge Haplotigs to generate a coverage histogram of the mapped PacBio reads. We set low (-l), middle (-m), and high (-h) in the coverage distribution of 10, 140, 195, respectively, and ran the second step in the Purge Haplotigs pipeline, ‘contigcov’, to analyze coverage on a contig-by-contig basis. Finally, we purged duplicate contigs using the function ‘purge’ to create a de-duplicated assembly. The resulting assembly (v2) was substantially smaller at 679 Mb with 84.5% fewer duplicated genes (*n* = 183).

We then polished the v2 assembly using our raw PacBio reads with Arrow, an algorithm in the variantCaller program within the Falcon software (Chin et al. 2013, 2016). Since this approach is not deterministic (i.e., repeated runs will produce slightly different results), we performed two rounds of polishing with Arrow (resulting in assemblies v3 and v4). For each round, we first re-mapped our raw PacBio reads to the latest assembly version (e.g., to make v3, we mapped raw PacBio reads to the v2 assembly) with Minimap2 v2.14-r883 (Li 2018) and the same flags as above. Next, we converted the BAM file of mapped reads to PacBio format with ‘pbbamify’ (part of the PacBio toolkit) with default flags. We sorted the re-formatted BAM file with samtools v1.9 as above (Li et al. 2009). We created an index of the mapped, sorted BAM with another PacBio tool, ‘pbindex’.

### Transcriptome assembly

To aid in genome annotation, we assembled an *O. amberensis* liver transcriptome from raw RNA-seq reads with Trinity v2.6.6 (Grabherr et al. 2011) using default settings and read trimming with Trimmomatic (Bolger et al. 2014) with flag “--trimmomatic.” We assessed transcriptome assembly completeness with BUSCO v3 and the 4,584 “actinopterygii_odb9” reference gene set (Simao et al. 2015).

### Genome annotation

We annotated the *O. amberensis* genome assembly (v4) using Maker2 v31.10 (Holt & Yandell, 2011). To support the annotation pipeline, we generated a repeat library using RepeatModeler v1.0.11 with default settings (A. F. Smit & Hubley, 2008). We also included gene evidence from the *O. amberensis* transcriptome and proteins for two other species, three-spined stickleback (*Gasterosteus aculeatus*) and Atlantic cod (*Gadus morhua*), which were both downloaded from Ensembl release 97 (http://ftp.ensembl.org/pub/release-97/). We first performed a *de novo* annotation of the *O. amberensis* genome assembly with our transcriptome and protein evidence with options est2genome = 1 and protein2genome = 1 in the Maker2 control file. Next, we used the gene finding program, SNAP (Korf, 2004), to generate *ab initio* gene predictions for our assembly. We first merged all of our gene models from our *de novo* Maker2 together into a single GFF3 using the Maker2 tool “gff3_merge” and converted them to ZFF format with “maker2zff.” We trained SNAP on these data by first breaking up our sequences into one gene per locus (with 1000 bp on either side of the predicted gene) using “fathom” (part of SNAP). We estimated parameters for our genes using “forge” (also included with SNAP) before building a Hidden Markov Model for our annotations with the script “hmm-assembler.pl”. After SNAP training, we re-ran Maker2 with our trained SNAP models by adding them in the control file (“snaphmm” option) and setting est2genome and protein2genome both to 0. We performed a second round of SNAP training using the same approach followed by running Maker2 for a third time. Next, we prepared to train another gene predictor, Augustus v2.5.5 (Stanke & Waack, 2003), on our gene models by first merging the models produced in the second round the SNAP training using “gff3_merge”, converting them to ZFF with “maker2zff”, and then to GBK format with a custom perl script (https://github.com/hyphaltip/genome-scripts/blob/master/gene_prediction/zff2augustus_gbk.pl). As Augustus is a machine learning approach, we first split our data set into a training and test set using the script “randomSplit.pl.” We then created a new “species” for Augustus with the script “new_species.pl” then performed training by running the “etraining” function on our training data set followed by evaluating its performance with the “augustus” tool. We optimized our prediction parameters with the script “optimize_augustus.pl.” After Augustus training, we re-ran Maker2 as above with the addition of the control file option “augustus_species.” Finally, we ran GeneMark for the *O. amberensis* genome assembly with the self-training algorithm using flag -ES. We then performed a final run of Maker2 with all of the gene model evidence included by adding the path to our GeneMark trained predictions using “gmhmm” to our control file along with the above evidence for Augustus (“augustus_species”) and SNAP (“snaphmm”). We kept est2genome and protein2genome as 0 for this final run.

### Repetitive element classification and annotation

We identified and classified repetitive elements (REs) in the genome assembly of *O. amberensis* using RepeatModeler v2.0.1 (Flynn et al., 2020). We then used the “queryRepeatDatabase.pl” script included with RepeatMasker v4.1.0 (A. Smit, Hubley, & Green, 2015) to output RepeatMasker’s internal library for ray-finned fishes (-species “actinopterygii”) and merged it with the *O. amberensis* custom repeat library generated by RepeatModeler. We used this combined Actinopterygii + *O. amberensis* library to annotate REs in the *O. amberensis* assembly using RepeatMasker v4.1.0 with the search engine set to “ncbi” and the -xsmall option. We repeated the above steps to classify and annotate REs in nine additional fish genome assemblies— *Chaenocephalus aceratus* (Chaenocephalus aceratus V1.0; Kim et al., 2019), *Cottoperca gobio* (Cottoperca gobio V1.0; Bista et al., 2020*), Danio rerio* (Danio rerio V4.0; Howe et al., 2013), *Dissostichus mawsoni* (Dissostichus mawsoni V1.0; Lee et al., 2021), *Gadus morhua* (gadMor1, Ensembl release 97; Star et al., 2011), *Gasterosteus aculeatus* (BROAD S1, Ensembl release 97; Jones et al., 2012), *Notothenia coriiceps* (Notothenia coriiceps V1.0; Shin et al., 2014), *Parachaenicthys charcoti* (Parachaenichthys charcoti V1.0, GigaDB dataset #100321; Ahn et al., 2017), and *Takifugu rubripes* (FUGU5, Ensembl release 98; Aparicio et al., 2002)—such that each assembly was annotated using a combined library of Actinopterygii and the custom library from RepeatModeler for each species. We summarized repeat abundance by parsing output from RepeatMasker and generating plots in R v3.5.1 (R Core Team, 2021) using custom scripts. To investigate patterns of transposable element (TE) activity through time we generated TE copy number divergence plots. We calculated Kimura substitution level (CpG adjusted) for transposable elements using the RepeatMasker utility script “calcDivergenceFromAlign.pl”, removed non-transposable element and unclassified repeats from the resulting “.divisum” file, and generated age distribution landscapes using RepeatMasker’s “createRepeatLandscape.pl” script. These plots are generated by comparing each TE copy to a reference sequence for that element. TE copies are placed along the x-axis based on their divergence relative to the consensus. Copy number peaks distributed on the left of the x-axis indicate abundant elements showing minimal divergence from the consensus and are considered indicative of recent TE activity, whereas right-distributed peaks indicate elements with high sequence divergence and more ancient TE activity.

### Selection on protein-coding genes

To assess selection on protein-coding genes in the *O. amberensis* genome, we calculated the ratio of non-synonymous to synonymous substitution (*d*_*N*_*/d*_*S*_) using CodeML with the PAML package (Yang, 2007). Coding sequences (CDS) for eight of the same genome assemblies as the repeat analyses above were included: *C. aceratus, C. gobio, D. rerio, G. aculeatus, G. morhua, N. coriiceps, O. amberensis*, and *T. rubripes*. For each species, we extracted the longest CDS for each gene from its annotation (GFF) file using AGAT v.0.7.0 (Dainat, 2020), first to retain only the longest isoform per gene (agat_sp_keep_longest_isoform.pl) and then write those isoforms to a FASTA file (agat_sp_extract_sequences.pl). We then translated CDS to protein using the function “transeq” in EMBOSS v.6.6.0 (Rice, Longden, & Bleasby, 2000). We identified orthologous groups of single-copy coding sequences using the OrthoVenn2 web server (L. Xu et al., 2019). We extracted corresponding CDS for each group of single-copy orthologs for each taxon. We aligned each of the orthologous CDS groups with PRANK v.170427 (Löytynoja, 2014) under a codon model. Before performing *d*_*N*_*/d*_*S*_ analyses, we generated a species tree with RAxML v.8.2.12 (Stamatakis, 2014) for a concatenated alignment of all of the single-copy orthologous nucleotide sequences with 100 bootstrap replicates using a GTR+Gamma model.

Using CodeML, we calculated *d*_*N*_*/d*_*S*_ for aligned single-copy orthologous coding sequences with branch-site models A1 (model = 2, NSsites = 2, fix_omega = 1) and A (model = 2, NSsites = 2, fix_omega = 0) and F3×4 codon frequency and cleandata = 1 for both. Using the same models, we varied the foreground lineage(s) by first focusing only on *O. amberensis* with all other lineages as background and second with the two notothenioids (*C. aceratus* and *P. charcoti*) as foreground lineages. We performed a Chi-Square Test with the pchisq() function in R v.3.6.3 (R Core Team, 2021) to compare our test model (A) to our null neutral model (A1) for each ortholog group. To control for multiple tests, we performed false discovery rate (FDR) correction in R using the p.adjust() function and the “Benjamini-Hochberg” method (Benjamini & Hochberg, 1995) with a cutoff of 0.05.

### Annotating hemoglobin and AFP regions

To explore the evolution of hemoglobin genes in *O. amberensis* and across the suborder Zoarcoidei, hemoglobin gene clusters were analyzed and compared to other species representing major perciformes lineages: *Cebidichthys violaceus* (Cvio_1.0, Heras, Chakraborty, Emerson, & German, 2020), *Anarrhichthys ocellatus* (GSC_Weel_1.0), *Pholis gunnellus* (fPhoGun1.1), *Perca flavescens* (PFLA_1.0, Feron et al., 2020), *Etheostoma spectabile* (UIUC_Espe_1.0, R. L. Moran, Catchen, & Fuller, 2020), *G. aculeatus* (BROAD S1), *Pseudoliparis sp*. (ASM433547v1, Mu et al., 2021), *Cottoperca gobio* (fCotGob3.1, Bista et al., 2020), *Ophiodon elongatus* (NWFSC_Oelon_r3.mflye, Longo et al., 2020), *Cyclopterus lumpus* (fCycLum1.pri, Holborn et al., 2021), and *Taurus bubalis* (fTauBub2.1). In teleost fish, hemoglobin genes occur in two distinct, unlinked clusters (LA and MN) which were first located in each assembly by searching for flanking genes (i.e., *rhbdf1b* and *aqp8*.*2* for the LA cluster and *kank2* and *nprl3* for the MN cluster). Sequence, length, position, and strand of individual exons of each alpha and beta gene were manually retrieved from each assembly by performing BLASTN v.2.12.0 searches and using the yellow perch *P. flavescens* as a reference for hemoglobin gene exon boundaries. Exons of each gene were concatenated and translated into protein sequence to verify their accuracy. If a gene was incomplete (e.g., a missing exon) or displayed a premature stop codon, the gene was considered pseudogenized.

To explore the evolution of AFPs across the Zoarcoidei (and an outgroup, *G. aculeatus*), regions associated with *afpIII* genes and their progenitor in eelpouts, sialic acid synthase (*nans*; Deng et al., 2010), were annotated by hand via a combination of BLASTN searches and targeted alignments. We focused on six species (five from Zoarcoidei) with high-quality genomic resources for the regions—five genome assemblies [*G. aculeatus* (BROAD S1), *O. amberensis, C. violaceous* (Cvio_1.0), *A. ocellatus* (GSCB_Weel_1.0), *P. gunnellus* (fPhoGun1.1)] and one set of bacterial artificial chromosomes (BACs) which targeted the same regions in another Antarctic eelpout, *Lycodicthys dearborni* (Deng et al., 2010). For non-*afpIII* genes (i.e., *nans* and all flanking genes), we used all known exons for *G. aculeatus* from Ensembl 104. For *afpIII* genes, we used a combination of the two exons described by Deng et al. (2010) for *L. dearborni* and Hobbs et al. (2020) for *P. gunnellus*. We first performed a BLASTN v.2.2.31 search with default settings against a reference database (created with “makeblastdb” for each genome or BAC group) and each exon as a separate query. We further investigated regions of interest by extracting them from our reference genome assemblies with samtools (H. Li et al., 2009) using the function “faidx” and aligning sequences with Clustal Omega (Sievers et al., 2011). To directly compare the utility of long-reads for studying *afpIII* genes to short reads, we assembled a genome from our DNA-seq data set using SPAdes v.3.15.3 (Bankevich et al., 2012) with default options. We then performed the same BLASTN v.2.2.31 search for *afpIII* genes described above.

Where appropriate, we assessed the quality of the assembly at target regions within the hemoglobin and AFP loci by mapping whole-genome, short-read Illumina data from a different *O. amberensis* individual to the *O. amberensis* PacBio genome assembly using bwa v.0.7.12 (Heng Li, 2013). We indexed the PacBio assembly using the “index” function then mapped reads with the “bwa-mem” algorithm with default settings. We converted the output from BAM to SAM format with samtools (H. Li et al., 2009) and visualized regions with the Integrated Genomics Viewer (Robinson et al., 2011). We used these data to assess coverage, presence or absence of indels, gaps, and any other features that could indicate a poor-quality assembly or error.

### Gene family expansion and contraction

To assess how gene families have changed along the *O. amberensis* lineage, we performed a statistical test of gene family expansion and contraction with CAFE v4.2.1 (De Bie, Cristianini, Demuth, & Hahn, 2006). For this analysis, we used the same eight fishes included in our *d*_*N*_/*d*_*S*_ analysis above, as well as the RAxML generated species tree and OrthoVenn2 gene orthology assignments. However, because CAFE requires an ultrametric species tree, we converted our species tree to ultrametric with r8s v1.81 (Sanderson, 2003) with a calibration point of 72.5 million years dating the split between *O. amberensis* and *G. aculeatus* (Betancur-R et al., 2013). For the CAFE analysis, we specified significance—i.e., the level at which a gene family change is considered rapid—at *P* < 0.01. We visualized gene ontology terms for rapidly evolving families using the online server of REVIGO (http://revigo.irb.hr/; Supek, Bošnjak, Škunca, & Šmuc, 2011).

## Results

### Sequence data

We generated 5,536,887 long-read PacBio sequences (65.7 Gb, mean length = 13.2 Kb, read N50 = 20 kb), 226,420,980 short-read DNA-seq reads, and 10,677,996 short-read RNA-seq reads. We estimated the *O. amberensis* genome to be 689.8 Mb from the DNA-seq data. Based on this estimate, our sequencing coverage for the PacBio data was 95x and 49x for the DNA-seq data. We estimated the genome heterozygosity to be 0.36% based on the DNA-seq data. The genome assembly, raw sequence data, and associated files for *O. amberensis* are deposited under NCBI BioProject PRJNA701078.

### O. amberensis genome assembly and annotation

Our final assembly of the *O. amberensis* genome (v4) was 680.7 Mb in 1,828 scaffolds with a scaffold N50 of 1.00 Mb. The longest scaffold was 6.98 Mb. We note that here and throughout this study, we use scaffold and contig interchangeably for *O. amberensis* so that we can use the same language for all of the assemblies included in this study. No scaffolding was performed on the *O. amberensis* assembly. Using a k-mer based estimate of genome size, our assembly represents 98.7% of the genome. The assembly contains 95.7% of the single-copy, conserved orthologous genes (BUSCOs) in the 4,584 gene reference set for Actinopterygii (Fig. 1D). With the same BUSCO gene set used for assessing the genome assembly completeness, the *O. amberensis* liver transcriptome used for annotation was 67.1% complete with 9.1% of genes fragmented and 23.8% missing. We annotated 22,572 genes in the *O. amberensis* genome assembly.

### Repetitive element content

Repetitive elements (REs) comprised 30.8% (209.6 Mb) of the *O. amberensis* genome assembly (Fig. 2). Of all repetitive sequences masked by RepeatMasker, 37.6% (78.8 Mb) were classified among the known types of repeats. DNA transposons accounted for the largest fraction of classified REs (33.8Mb or 16.2% of total REs), followed by long-interspersed nuclear elements (LINEs; 25.2Mb or 12.0%), long-terminal repeats (LTRs; 15.0 Mb or 7.2%), satellite DNAs (2.4 Mb or 1.1%), small interspersed nuclear elements (SINEs; 0.18 Mb, 0.1%) and other repeats (2.0 Mb, 1.0%). Unclassified REs accounted for the remaining repetitive component (130.1 Mb or 62.4% of total REs) of the *O. amberensis* assembly.

**Figure 2.**
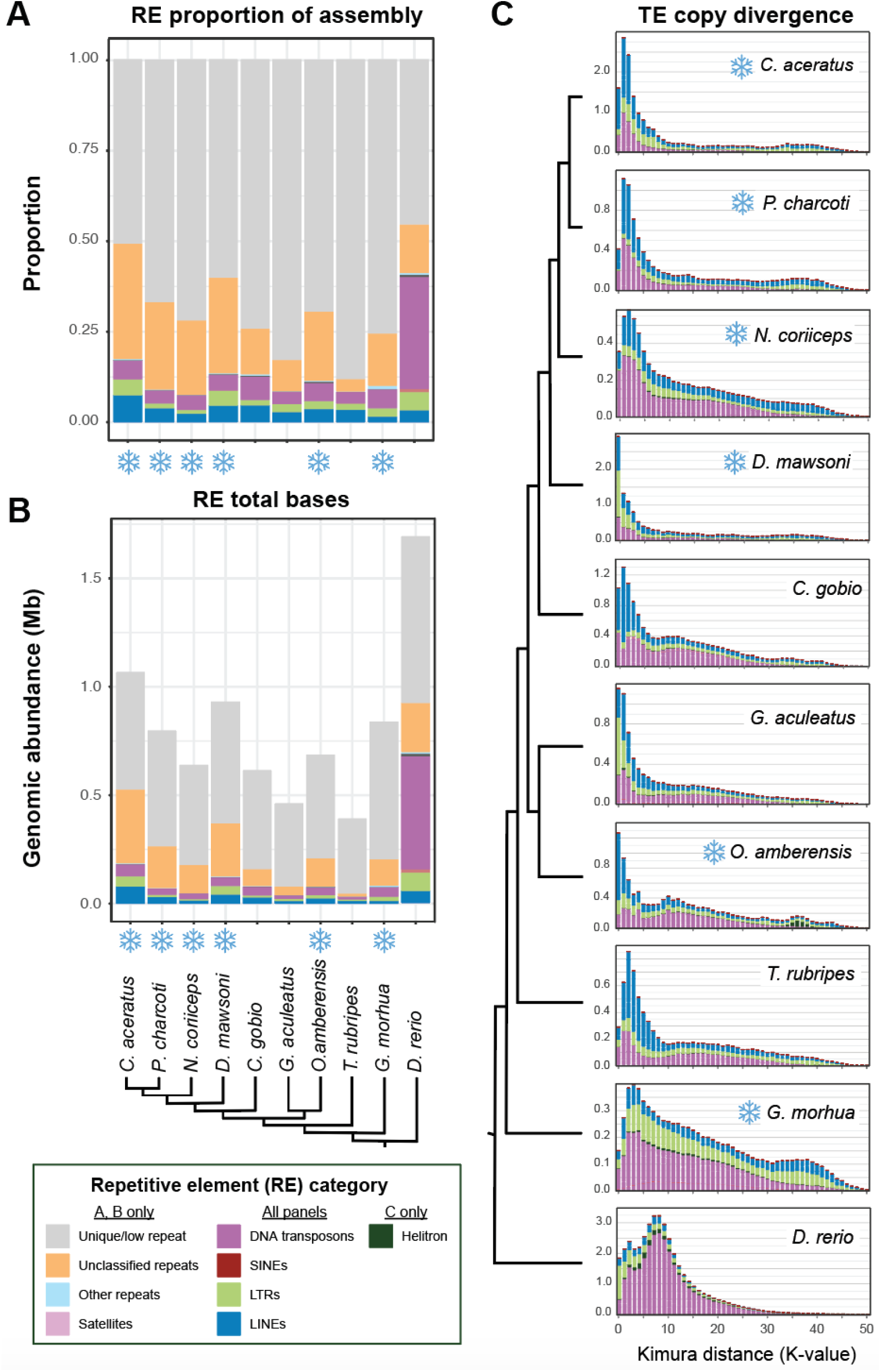
Repetitive element abundance in focal fishes. In all panels, snowflakes indicate species with distributions that include polar seas. (A) Abundance of major repetitive element (RE) categories shown as genomic proportions. (B) RE abundance shown as total bases in the assembly such that the height of each bar is proportional to the assembly length for that species, with the assembly fraction not masked by RepeatMasker shown in gray. (C) Transposable element (TE) copy divergence plots. The y-axis shows TE abundance as a proportion of the genome (e.g., 1.0 = 1% of the genome). The x-axis shows sequence divergence (CpG adjusted Kimura distance) relative to consensus sequences for TE superfamilies. Copy number peaks with abundance skewed toward the left show less sequence divergence from the consensus and potentially correspond to TE copies with a recent history of activity relative to peaks with right-skewed abundance which potentially correspond to remnants of more ancient TE copies. For visual display, TE superfamilies are grouped by DNA transposons, Helitrons, LTRs, LINEs, and SINEs and colored accordingly. A similar plot with separated superfamilies is shown in Fig. S1.

*Opthalmolycus amberensis* and the four Antarctic notothenioid species (*C. aceratus, D. mawsoni, N. coriiceps*, and *P. charcoti*) showed greater RE abundance in both genomic proportion and total repeats (genomic proportion and total repeats = 28.2% and 179.5 Mb; 30.8% and 209.6 Mb; 33.3% and 265.0 Mb; 40.1% and 371.4 Mb; 49.4% and 526.5 Mb in *N. coriiceps, O. amberensis, P. charcoti, D. mawsoni*, and *C. aceratus*, respectively) compared to non-Antarctic ingroup species. Arctic-adapted *G. morhua* (24.6% and 205.0 Mb) and the non-polar sister taxon to the notothenoids, *C. gobio* (26.1% and 159.0 Mb), had the next greatest RE abundance among ingroup taxa, while *T. rubripes* (11.9% and 46.7 Mb) and *G. aculeatus* (17.2% and 79.5 Mb) have smaller, less repetitive genome assemblies. Although satellite DNAs make up a small fraction of REs in all species, satellites were more abundant in the *O. amberensis* assembly compared to all other in-group species. For all species either DNA transposons or LINEs comprised the largest proportion of classified repeats (Fig. 2).

Transposable element (TE) copy divergence plots in *O. amberensis* suggested evidence of dynamic shifts in transposable activity through time (Fig. 2). LINEs dominated recent and ongoing TE activity (Fig. 2C), which was driven by recent expansion of *L2* and *Rex/Babar* elements, accounting for much of the LINE abundance (blue peak between *K* = 0–5; Fig. 2C, Fig. S1). DNA transposons showed a relatively older history of proliferation dominated by *hAT* elements (broad peak centered at *K* = 10; Fig. 2C, Fig. S1), though some more recent peaks are also evident. Prior to this putative increase in DNA transposon activity, Helitrons showed evidence of an ancient burst with abundant sequences centered over *K =* 35–40 (Fig. 2C, Fig. S1). LTRs appear to have maintained modest activity through time (Fig. 2C), although *ERV1* and *Gypsy* elements show some evidence of recent expansion (Fig. S1). Similar trends of recent LINE expansions were visible in the other two Antarctic species, *C. aceratus* and *P. charcoti*, as well as *T. rubripes*. Recent DNA transposon activity was evident in all species, though least pronounced in *O. amberensis. Gasterosteus aculeatus* and *D. mawsoni* were the only species for which LTRs dominated recent TE activity (mostly *Gypsy, ERV1, DIRS*, and *Pao* elements; Fig. 2C, Fig. S1). Three other polar species, *C. aceratus, P. charcoti*, and *G. morhua*, all had evidence for ancient TE expansions similar to the Helitron expansion in *O. amberensis*, however in these species LINEs and LTRs drove the expansion (Fig. 2C, Fig. S1).

### Selection on protein-coding genes

We identified 4,726 single-copy orthologous genes that were present in all eight species. When only *O. amberensis* was the foreground lineage in *d*_*N*_*/d*_*S*_ analyses, we identified 229 protein-coding genes with evidence of positive selection with FDR < 0.05 (Fig. 3). When the two notothenioids (*C. aceratus* and *N. coriiceps*) were the foreground lineages, 371 protein-coding genes exhibited evidence of positive selection. When these two lists were intersected, just 13 protein-coding genes exhibited evidence of positive selection in both groups, leaving 216 (94.3%) and 358 (96.5%) genes that were uniquely under positive selection in *O. amberensis* or the notothenioids, respectively (Fig. 3). The gene that exhibited the strongest signature of positive selection in *O. amberensis* was *bach2b* (Ensembl ID #ENSGACG00000006245; FDR = 5.4e-11) which has been linked to embryonic development in zebrafish (Cohen et al., 2020). For the notothenioids, the strongest signature of positive selection was *eif4enif1* (FDR = 6.03e-20), a eukaryotic translation initiation factor which has been linked to thermal stress in salmonids (Akbarzadeh et al., 2018).

**Figure 3.**
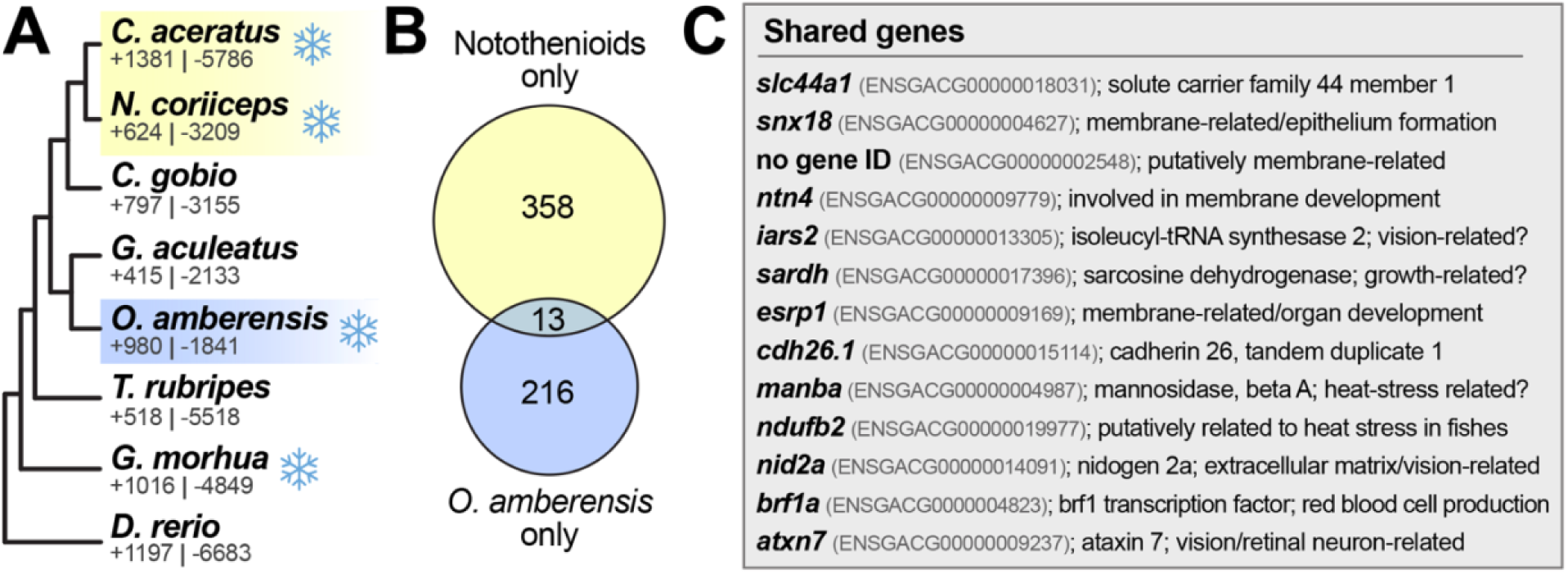
(A) The maximum-likelihood guide tree generated for *d*_*N*_*/d*_*S*_ analyses from 4,726 protein-coding genes. Yellow and blue boxes indicate the foreground lineages in respective analyses (top, yellow: notothenioids only; bottom, blue: *O. amberensis* only). Snowflakes indicate species with distributions that include polar seas (Antarctic: *C. aceratus, N. coriiceps, O. amberensis*; Arctic: *G. morhua*). Numbers under species names indicate the number of gene families that were expanded (+) or contracted (-) in each lineage. (B) The number of genes with evidence of positive selection in each independent analysis and in both analyses (middle of the Venn diagram). (C) The 13 genes that were shared between both analyses. When applicable, gene names and IDs from the ortholog in stickleback (*Gasterosteus aculeatus*, BROAD S1) are given. The FDR cutoff used in all analyses was 0.05.

### Hemoglobin and AFPs

In *O. amberensis* and across Zoarcoidei, the hemoglobin LA cluster has remained stable with the typical organization of two alpha-globin and one beta-globin genes (Fig. 4). However, among perciformes, the beta-gene of the LA cluster, *hbbla*, was repeatedly lost or pseudogenized in some species, including in the Zorcoidei monkeyface prickleback *Cebidichthys violaceus* (Fig. 4b). In contrast, the MN cluster in perciformes underwent dynamic changes in a lineage specific fashion and harbored up to 13 alpha-globin and 15 beta-globin genes in the Longspined bullhead sculpin (*Taurulus bubalis*) and numerous instances of pseudogenization, especially frequent in the Orangethroat darter *Etheostoma spectabile* (Fig. 4c). The MN cluster organization could not be analyzed in the wolf-eel *A. ocellatus* due to the lack of contiguity in the assembly at this locus. Within Zoarcoidei, however, the MN cluster composition remained relatively stable among the three studied species with three alpha-globin genes and two or three functional beta-genes. A unique feature of the Zoarcoidei relative to the other studied species is the presence of a *hbbmn2* pseudogene in *C. violaceus* and *P. gunnellus* and the loss of this gene in *O. amberensis* (Fig. 4c).

**Figure 4.**
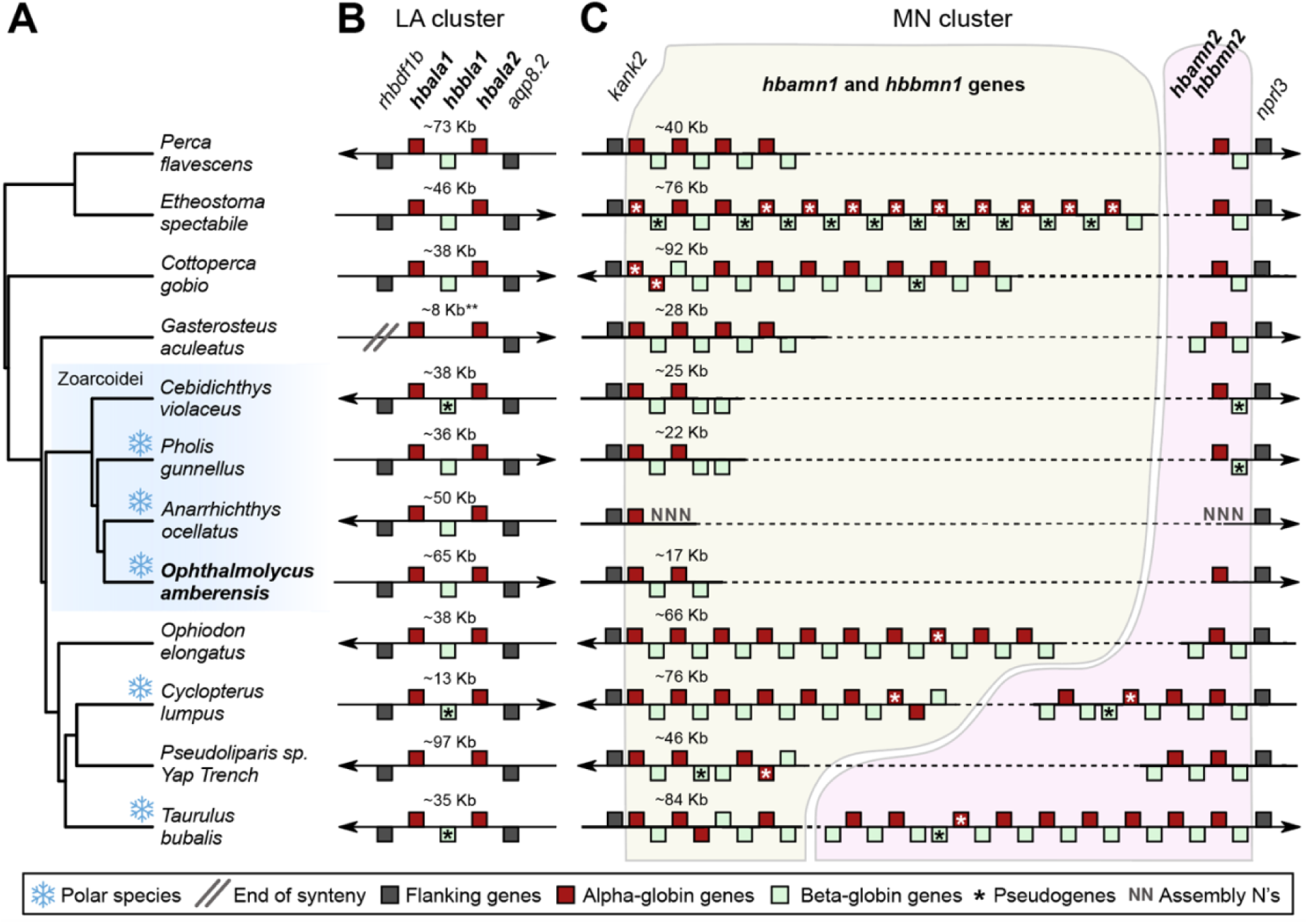
Evolution of the hemoglobin clusters in Perciformes. (A) The phylogenetic relationship of the studied perciform species was extracted from Rabosky et al. (2018). (B) Genomic organization of the hemoglobin LA cluster flanked by the genes *rhbdf1b* and *aqp8*.*2* in all studied species. (C) Genomic organization of the hemoglobin MN cluster flanked by the genes *kank2* and *nprl3* in all studied species. Genes flanking the clusters are represented by dark-grey squares, alpha-globin genes (*hba*) are represented with red squares, beta-globin genes (*hbb*) are represented with light green squares, and pseudogenes are labelled with an asterisk (*). The arrowheads indicate the direction of the clusters in each assembly. The position of each square above or below the line denotes the position of the gene on the forward (right arrow) or reverse strand (left arrow), respectively. “N” in *A. ocellatus* denote that the assembly contains gaps between exon1 of *hbamn1* and *nprl3*, therefore preventing the study of this cluster in this species. Approximate lengths of the regions are indicated over the respective clusters. Basic metrics for the genome assemblies included in this analysis are provided in Table S1. Complete annotations for the LA and MN clusters are provided in Tables S2 and S3, respectively.

For the *afpIII* region in Zoarcoidei between *sncgb* and *ldb3b*, we found no evidence of *afpIII* genes in the outgroup, *G. aculeatus*, nor in one of our in-group species, *C. violaceus* (family Stichaeidae; Fig. 5). Among the four species with copies of some portion of an *afpIII* gene (either exon 1, exon 2, or complete copies of both exons 1 and 2 in order), copy number varied widely. *Lycodicthys dearborni* has at least 23 complete copies of *afpIII* with some copies of exon 1 in isolation scattered throughout the same region (Fig. 5). *Lycodicthys dearborni* also has two psuedogenized *afpIII* copies (Fig. 5). Similar to *L. dearborni, O. amberensis* also had a high number of complete *afpIII copies* (at least 13) with some copies of exon 1 in the same region. *Pholis gunnellus* has at least 15 complete *afpIII* copies as well as partial copies. Finally, *A. ocellatus* has evidence of *afpIII* genes but with four complete *afpIII* copies spread over four of the six scaffolds including components of the *afpIII* gene, it is unclear how many copies are truly present (Fig. 5). This is primarily because, unlike *L. dearborni* for instance, no *A. ocellatus* scaffold has evidence of both *afpIII* genes and flanking regions. With our short-read, DNA-seq assembly for *O. amberensis*, we confirmed the useful but coarse nature of inference stemming from short-read data for the study of complex genomic regions. Indeed, similar to *A. ocellatus*, we found evidence for five partial copies of *afpIII* spread across five different scaffolds. The sialic acid synthase (*nans*) region—the progenitor of *afpIII* in eelpouts (Deng et al., 2010)—is generally well-conserved across and beyond Zoarcoidei with only one missing copy in *P. gunnellus* (*nansb*; Fig. S2).

**Figure 5.**
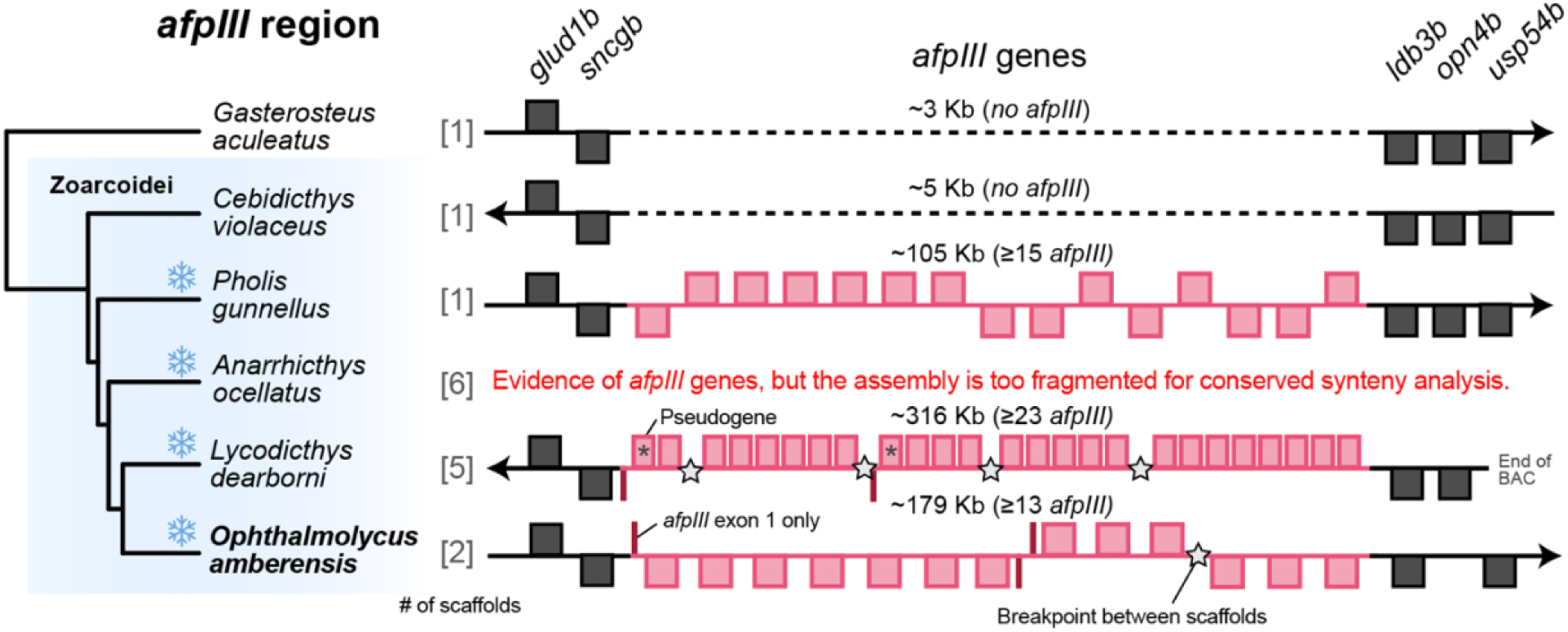
Synteny *afpIII* region in the suborder Zoarcoidei with three-spined stickleback (*G. aculeatus*) as the outgroup. Snowflakes indicate species with distributions that overlap polar regions and stars indicate breakpoints between scaffolds. As in Fig. 4, the arrowheads indicate the direction of the clusters in each assembly. The position of each square above or below the line denotes the position of the gene on the forward (when the arrow points to the right) or reverse strand (when the arrow points to the left), respectively. Basic metrics for the genome assemblies included in this analysis (all species except for *L. dearborni* which is not a genome assembly) are provided in Table S1. Complete annotations for all *afpIII* copies are provided in Table S4. The phylogenetic relationships shown were extracted from Hotaling, Borowiec, et al. (2021) and Betancur-R et al. (2013).

### Gene family expansion and contraction

Along the *O. amberensis* lineage (and after its divergence from *G. aculeatus*), we identified 980 gene families that have expanded and 1,841 that have contracted with 137 rapidly changing families at *P* < 0.01 (110 expanded, 27 contracted; Fig. 3A, Table S5). Relative to the other species included, the number of expanded gene families for *O. amberensis* was intermediate while the total number of contracted families was the lowest (Fig. 3A). Gene ontology terms for rapidly expanding and contracting gene families encompassed a wide array of biological processes including genes linked to immune system response, vision, membrane development, and DNA repair (Figs. S3-S4).

## Discussion

The rapid accumulation of genomic resources across the tree of life is empowering a new era of comparative genome science (Feng et al., 2020; Hoencamp et al., 2021; Hotaling, Kelley, et al., 2021; Marks, Hotaling, Frandsen, & VanBuren, 2021; Thomas et al., 2020). Moreover, the rise of long-read sequencing has improved the contiguity of difficult to assemble, repeat-rich portions of genomes. Indeed, genome assemblies that integrate long-reads are on average 48x more contiguous than those that do not (Hotaling, Sproul, et al., 2021). In this study, we generated and compared a new, high-quality genome assembly for *O. amberensis* to other polar and non-polar fishes with an emphasis on aspects that are difficult or impossible to assess with short-reads alone (repetitive elements and repeat-rich, dynamic loci). Collectively, our results highlight the power of long reads for resolving complex, repetitive genomic regions of interest and provide new understanding of adaptation to the extreme conditions of the Southern Ocean.

### Repetitive element dynamics

Repetitive elements (REs) are major drivers of genome evolution in fish and their abundance varies widely across species. Transposable elements (TEs) make up the largest fraction of REs in fish genomes ranging from 5–56% (Shao, Han, & Peng, 2019). *Ophthalmolycus amberensis* is intermediate among fish lineages with classified TEs comprising nearly 11% of the assembly and having an overall RE genomic proportion of ∼31%. Our findings generally corroborate previous studies that show DNA transposons and LINEs are consistently abundant in fish genomes (e.g., Chalopin, Naville, Plard, Galiana, & Volff, 2015) and that RE abundance roughly correlates with genome size. The age distribution of TEs in *O. amberensis* revealed dynamic shifts in its TE landscape. While LINEs dominated recent TE activity, DNA transposons and Helitrons both showed earlier bursts of activity.

Recent studies have highlighted a potential link between environmental stress and bursts of TE activity in fish (Chénais, Caruso, Hiard, & Casse, 2012). In Antarctic notothenioid fish, *DIRS1* retrotransposons form hotspots of chromosome rearrangements within lineages, potentially driving speciation (Auvinet et al., 2018; Auvinet et al., 2019). Consistent with this, we found that *DIRS* elements show high levels of recent activity in notothenioid species *C. aceratus*, and *D. mawsoni*, but not in other polar fish lineages, highlighting the dynamic evolution of these elements in notothenoids (Fig. S1). Heating and cooling cycles in polar seas have also been hypothesized to induce environmental stress which in turn drives bursts of TE activity (Auvinet et al., 2018). We found that across polar-adapted lineages, four of six species (*O. amberensis, C. aceratus, P. charcoti*, and *G. morhua*) exhibited evidence consistent with ancient TE bursts of similar relative timing (i.e., similar in timing to the peak of Helitron activity in *O. amberensis*). This modest, but distinct pattern is observed independently in three lineages that lack of phylogenetic (and spatial) overlap (Fig. 2, Fig. S1). While it is premature to assume that these signatures reflect historical, climate-induced shifts that affected fishes at both poles, we view this pattern an interesting preliminary finding that merits additional investigation. Future efforts to integrate dense taxonomic sampling will allow for contextualizing links between TE activity, environmental stress, and genome evolution, particularly as they relate to ancient and contemporary environmental change.

### Selection on protein-coding genes

Across thousands of shared orthologs, only a small fraction exhibited signatures of selection in either polar fish group (notothenioids or *O. amberensis*), and of those, just 13 genes were shared between the two. The split between Zoarcoidei and Notothenioidei occurred roughly 80 million years ago (Rabosky et al., 2018). Thus, with a moderate amount of shared ancestry, if adaptation to cold environments relies on universal biological and physiological mechanisms, we might expect a signature of evolution in both groups with significant overlap between the targets of selection. This was not the case. However, among the overlap, three themes emerged: selection has acted on genes involved in (i) membrane structure, (ii) thermal stress related pathways, and (iii) vision.

Membranes play an obvious role in organismal and cell survival as they delimit cellular structures, mediate transmembrane movement of solutes and other materials, and play roles in other key processes and structures (Rothfield, 2014). Moreover, because environmental conditions often play a role in determining how lipids—a component of biological membranes— arrange themselves (Rothfield, 2014), it is unsurprising that membranes are important for thermal adaptation in a wide variety of organisms (Hazel, 1995). Fishes living in the most consistently cold marine habitats on Earth have experienced positive selection on genes related to membrane structure and function (e.g., *esrp1, ntn4, snx18*). Generally speaking, membrane fluidity is maintained as temperature changes, a phenomenon known as homeoviscous adaptation (Hazel, 1995). However, the extreme, but stable, cold temperatures of polar seas means that maintenance of a homeoviscous adaptation response to changing conditions should not hold any fitness benefit (Cossins & Prosser, 1978). Still, polar fish membranes must function at low temperature and directional selection may have acted on this trait. For instance, one target of selection in polar fishes, *snx18*, in concert with *fip5*, is involved in the morphogenesis of the epithelial lumen including the formation of membrane tubules (Willenborg et al., 2011).

While the exact role of selection on membrane-associated genes in polar fish remains to be determined, it is plausible that at least one driver is functioning in the cold.

The ability to mitigate thermal stress is a hallmark of polar adaptation and has been implicated in a variety of polar fish traits, most notably the evolution and expansion of AFPs. Less clear is how selection may have shaped the evolution of genes involved in thermal stress pathways. To this end, we identified two general thermal stress-related targets of selection in polar fishes: *manba* and *ndufb2*. Both have been observed to be downregulated under heat stress in oysters (Ding et al., 2020) and trout (Garvin, Thorgaard, & Narum, 2015), respectively, with potential implications for thermal plasticity (e.g., Kenkel & Matz, 2016). While plasticity cannot be assessed in our data set, nor do we expect thermal plasticity to be particularly important for fishes living in the consistently subfreezing Southern Ocean, links between these molecular targets and thermal tolerance in a wide range of organisms suggests that future work to assess their role in cold adaptation may be warranted.

Vision-associated gene family expansions or selections is commonly observed for cave-dwelling or deep sea fishes (e.g., Mu et al., 2021). All three of our focal Antarctic lineages—*O. amberensis, C. aceratus*, and *N. coriiceps*—can be found hundreds of meters below the surface. Moreover, the extreme nature of seasonal light at high latitudes and extended periods of sea ice cover could cause cave-like conditions for fishes and other marine taxa, regardless of where they reside in the water column. Highlighting the power of long-reads for genomic inquiry, we identified rapidly evolving gene families that were also linked to vision. For instance, three additional copies of *gja3* are present in *O. amberensis* relative to other species in the analysis and all five copies are arrayed in a ∼135-Kb region of the same scaffold (tig00024373). In humans, *GJA3* has been implicated in vision defects (Graw, 2003). In addition to gene copy expansions in vision-related genes, we also detected a shared signature of selection on vision-associated genes (*iars2, nid2a*, and *atxn7*) in two disparate lineages of polar fish. In mammals, *IARS2* has been associated with cataracts (Vona et al., 2018) and it has also been under selection in another ray-fined, mesopelagic fish *Lamprus guttatus* (Wang et al., 2022). The other two genes we identified have been recently linked to degeneration of retinal neurons (*atxn7*, Sun, Chen, & Jin, 2020) and eye development (*nid2a*, Carrara, Weaver, Piedade, Vöcking, & Famulski, 2019). Our capacity to assess selection on single copy orthologs and gene family expansions further emphasizes the utility of long reads for the study of adaptations that may be linked structural genomic change.

### Evolution of hemoglobin genes in Zoarcoidei

Hemoglobin is a major trait for adaptation to different environments in fishes (Powers, 1980; Verde, Giordano, di Prisco, & Andersen, 2012). Because the capacity of hemoglobin to bind oxygen is sensitive to environmental conditions such as temperature and pH, the molecular evolution of hemoglobin genes has been proposed to be necessary for fish to maintain respiratory homeostasis in new or changing environments (Powers, 1980). Vertebrate hemoglobin is a tetrameric protein composed of two alpha-globin and two beta-globin subunits. Combinations of different alpha and beta subunits can result in multiple hemoglobin tetramers with different functions. This “hemoglobin multiplicity” may also be a source of adaptive potential (Brix, Clements, & Wells, 1999; Houston & Gingras-Bedard, 1994; Verde et al., 2012). Hemoglobin multiplicity may be, at least in part, linked to hemoglobin gene copy number. For instance, cod fishes living in shallow and unstable environments have more hemoglobin gene copies than species living in more stable deep-sea environments (Baalsrud, Voje, et al., 2017). The Southern Ocean is a chronically cold and hyper-oxygenated environment, yet stable marine habitat. In this environment, the suborder Notothenioidei, which includes the white-blooded Antarctic icefishes lacking functional hemoglobin (Kim et al., 2019; Near et al., 2006), are the most common fish species. A reduction of hemoglobin multiplicity prior to cold adaptation followed by a decline in oxygen affinity has been documented in cold-adapted notothenioids (Verde, Parisi, & di Prisco, 2006). In contrast, studies of hemoglobin evolution in other polar species, as well as other Zoarcoidei, remain scarce (but see, di Prisco et al., 1990; Verde et al., 2002; Weber, Hourdez, Knowles, & Lallier, 2003). Eelpouts and other species in the suborder Zoarcodei, however, inhabit diverse environments from pole to pole and from the surface to hadal depths, making them excellent models for understanding hemoglobin evolution across disparate environments.

Across ray-finned fishes in the order Perciformes living in a wide range of habitats, the two teleost hemoglobin clusters, LA and MN (Opazo, Butts, Nery, Storz, & Hoffmann, 2013), have followed dramatically different evolutionary paths (Fig. 4). For the most part, the LA cluster has remained stable across species, with two alpha and one beta-globin genes (Opazo et al. 2013), and only a few instances of pseudogenization or loss of the beta-globin gene (Fig. 4a). In contrast, the MN cluster showed dramatic lineage-specific evolution, in line with other teleosts (Opazo et al., 2013). In most species in our comparison, the MN cluster expanded by multiple tandem duplications with frequent pseudogenization. In contrast, the genetic organization of the MN cluster has been relatively stable within Zoarcoidei, with three alpha-globin and two or three beta-globin genes (Fig. 4b). However, all studied Zoarcoidei share either the pseudogenization or loss of the *hbbmn2* gene, which is predicted to be functional in all other perciformes included. All studied Zoarcoidei appear to have three to five hemoglobins (di Prisco et al., 1990; Verde et al., 2002; Weber et al., 2003) despite a relatively low number of hemoglobin genes in comparison to other perciformes. Thus, at least for Zoarcoidei, globin gene copy number does not seem to correlate with hemoglobin multiplicity. The relative stability of the Zoarcoidei MN cluster is also remarkable as the studied species inhabit similar environments to other species with highly variable MN clusters. This suggests that adaptation in Zoarcoidei hemoglobin may occur at the gene sequence level rather than duplication as seen in these other perciformes. The study of hemoglobin should be pursued in more Zoarcoidei species, including in the globally distributed eelpout genera (Hotaling, Borowiec, et al., 2021), to better understand the molecular mechanisms underlying adaptation to environmental conditions.

### Evolution of antifreeze proteins

The evolution of antifreeze proteins (AFP) is a hallmark of adaptation to cold environments across a wide range of taxa (Atıcı & Nalbantoǧlu, 2003; Davies et al., 2002). Multiple routes to AFP evolution have been documented in polar fishes, including the evolution of *afpIII* from neofunctionalization of a duplicated sialic acid synthase b (*nansb*) gene in the Zoarcoidei (Deng et al., 2010) and the evolution of AFGPs in both notothenioids and Arctic cod (Chen et al., 1997a, 1997b; Zhuang et al., 2019) via two distinct molecular routes (Baalsrud, Tørresen, et al., 2017). Independent evolution of AFPs through non-overlapping pathways is one of the most well-known differences between the adaptation of eelpouts and notothenioids to the Southern Ocean. However, understanding of AFP evolution within the suborder Zoarcoidei—for which eelpouts comprise the most speciose family (Zoarcidae)—remains incomplete, largely due to the complexity of its history with many species and isoforms evolving at both poles and the complexity of the genomic region underlying it. Recent phylogenetic evidence suggests that *afpIII* arose only once, early in the evolution of the Zoarcoidei at least 18 million years ago (Hobbs et al., 2020). The lack of *afpIII* evidence in the typical region of the genome for Zoarcoidei for monkeyface prickleback *C. violaceus* (family Stichaeidae) poses an additional question about the evolution of AFPs in Zoarcoidei. To our understanding, every previously surveyed species in the suborder has exhibited *apfIII* sequence evidence (at least 14 species from six families, Hobbs et al., 2020). Given that *C. violaceus* is part of the Zoarcoidei “in group” (Hotaling, Borowiec, et al., 2021), its lack of similar *apfIII* evidence suggests that most likely the *afpIII* gene has been secondarily lost in *C. violaceus*. Other less likely possibilities are that the locus did not arise only once at the base of the clade or it has been horizontally transferred within the clade (e.g., Graham, Lougheed, Ewart, & Davies, 2008). It is also possible that this finding could be due to assembly error, but this is unlikely given the assembly’s high contiguity (scaffold N50 = 6.72 Mb, Table S1), lack of N’s in the predicted *afpIII* region, and contiguity of flanking regions (Fig. 5).

Gene duplication, rearrangement, and modification has long obscured the study of AFP evolution (e.g., Deng et al., 2010). *afpIII* copy number varies considerably between closely related species (Scott, Fletcher, & Davies, 1986) and even within species across their geographic range (Hew et al., 1988). Short-read genome assemblies limit *afpIII* copy number inference. For instance, a BLAST search of a short-read assembly for *O. amberensis* identified five partial *afpIII* copies spread across five different scaffolds. While there are more robust ways to detect gene copy numbers from short-read genome assemblies (e.g., through liberal BLAST searches and phylogenetics, Baalsrud, Tørresen, et al., 2017), the reality that genomic organization cannot be resolved remains. Long-read assemblies can largely ameliorate this issue. Indeed, following the massive improvements of long-read sequencing for genome assembly, a complete sequence of the complex *afpIII* locus has now been produced (for *P. gunnellus*) with a second, near complete sequence reported in this study. For both Antarctic eelpouts, *O. amberensis* and *L. dearborni*, and the rock gunnell, *P. gunnellus*, a large number of similar but not identical *afpIII* copies (at least 13, 23, and 15 respectively, Fig. 5) may represent an evolutionarily tuned system for depressing the serum freezing point (Kelley, Aagaard, MacCoss, & Swanson, 2010).

### Adaptation to the Southern Ocean: new insights from long-read sequencing

We have entered an exciting time for genome biology. It is now possible to characterize complex regions of genomes to better understand evolution across virtually any group of interest. In this study, we used long-read sequencing to generate a high-quality reference assembly for an Antarctic eelpout, *O. amberensis*, the first for the group. We used this new genomic resource to better understand the evolution of *O. amberensis* in the Southern Ocean, particularly as it relates to other Antarctic fish specialists (i.e., notothenioids) and the rest of the suborder Zoarcoidei. We showed evidence of historical shifts in TE activity in *O. amberensis* and other polar fishes, perhaps reflecting an ancient burst of TE activity in response to environmental change. Moreover, we found little overlap between the targets of natural selection in *O. amberensis* and in notothenioids inhabiting the same habitats, indicating that adaptation of both groups to the Southern Ocean appear to have followed largely different trajectories. However, we did find overlap in a few targets of selection, many of which were genes involved in vision or membrane structure. We also observed the relative evolutionary stability of the hemoglobin MN cluster in the Zoarcoidei relative to other teleosts despite these species inhabiting similar environments to species with much more diverse MN clusters. For AFPs, we identified the first species in the suborder Zoarcoidei with no evidence of *afpIII* genes in the genomic region where they occur for the group (*C. violaceus*), likely reflecting a lineage-specific loss of the cluster. To further understand the evolution of polar fishes, both within and across groups, more dense taxonomic sampling is needed. Snailfishes (order Scorpaeniformes; family Liparidae) occur around the world, including at hadal depths, and are the most speciose deep-water family in the Southern Ocean, making them another exceptional candidate for comparisons to understand the general rules of Antarctic adaptation.

Beyond polar fish adaptation, the utility of long-read sequencing for understanding complex regions of genomes is clear. The power of these data will increase as accuracy continues to improve (e.g., circular consensus sequencing (Wenger et al., 2019) and reads lengthen. The application of long-read technologies to eelpouts and other zoarcids will yield additional high-quality, contiguous sequences of the *afpIII* locus. The limiting factor for understanding questions similar to those posed in this study is shifting from generating high-quality assemblies to obtaining samples compatible with high-molecular weight DNA extraction and long-read sequencing. We echo previous studies (e.g., Buckner, Sanders, Faircloth, & Chakrabarty, 2021; Hotaling, Kelley, et al., 2021) that the field needs curated specimen vouchers with properly stored tissues and associated metadata to empower the future of genome biology in polar fishes and beyond.

## Acknowledgements

We thank the captain and crew of the ARSV *Laurence M. Gould*, the personnel of the U.S. Antarctic Program Support Contractors for assistance in Chile, at sea, and at Palmer Station, as well as the logistics in Denver, CO for their support related to the Antarctic fieldwork required for this study. We also thank Blair Perry, Kerry McGowan, and Ellie Armstrong for comments that improved the manuscript. We acknowledge funding from NSF awards OPP-1543383 and OPP-1947040 (supporting T.D.), and OPP-1906015 (to J.L.K.) as well as funding from the Antarctic Science Bursary. The RNA sequencing data used in this study was provided as part of a workshop sponsored by the NSF (OPP-1744877) and a Grant in Aid of Research through Long Island University-Post (both to Scott Santagata). We acknowledge computing resources from the Center for Institutional Research Computing at Washington State University and the NSF Extreme Science and Engineering Discovery Environment (XSEDE).

## Data accessibility

The genome assembly for *O. amberensis*, raw sequence data, and scripts used for analysis are deposited under NCBI BioProject #PRJNA701078 and this study’s GitHub (https://github.com/scotthotaling/oamberensis_genome).

## Author contributions

S.H. and J.L.K. conceived of the study. L.S.F.L. and S.H. performed laboratory work. S.H., T.D., J.S.S. and J.L.K. performed analyses. S.H. led the writing of the manuscript with considerable help from T.D., J.S.S., and J.L.K. T.D. provided the *O. amberensis* samples. All authors read and approved the final version of the manuscript.

## Notes

### Competing Interest Statement

The authors have declared no competing interest.

### Summary of Updates

Updated and expanded analyses. Refined figures.

## References

Ahn, D.-H., Shin, S. C., Kim, B.-M., Kang, S., Kim, J.-H., Ahn, I., … Park, H. (2017). Draft genome of the Antarctic dragonfish, Parachaenichthys charcoti. GigaScience, 6(8), gix060.

Akbarzadeh, A., Günther, O. P., Houde, A. L., Li, S., Ming, T. J., Jeffries, K. M., … Miller, K. M. (2018). Developing specific molecular biomarkers for thermal stress in salmonids. BMC Genomics, 19(1), 1–28.

Anderson, M. E. (1994). Systematics and osteology of the Zoarcidae (Teleostei: Perciformes). Ichthyological Bulletin of the J.L.B. Smith Institute of Ichthyology, 60, 1–120.

Andrews, S. (2010). FastQC: a quality control tool for high throughput sequence data.

Aparicio, S., Chapman, J., Stupka, E., Putnam, N., Chia, J.-m., Dehal, P., … Smit, A. (2002). Whole-genome shotgun assembly and analysis of the genome of Fugu rubripes. Science, 297(5585), 1301–1310.

Atici, Ö., & Nalbantoǧlu, B. (2003). Antifreeze proteins in higher plants. Phytochemistry, 64(7), 1187–1196.

Auvinet, J., Graça, P., Belkadi, L., Petit, L., Bonnivard, E., Dettaï, A., … Higuet, D. (2018). Mobilization of retrotransposons as a cause of chromosomal diversification and rapid speciation: the case for the Antarctic teleost genus Trematomus. BMC Genomics, 19(1), 1–18.

Auvinet, J., Graça, P., Ghigliotti, L., Pisano, E., Dettaï, A., Ozouf-Costaz, C., & Higuet, D. (2019). Insertion hot spots of DIRS1 Retrotransposon and chromosomal diversifications among the Antarctic Teleosts Nototheniidae. International journal of molecular sciences, 20(3), 701.

Baalsrud, H. T., Tørresen, O. K., Solbakken, M. H., Salzburger, W., Hanel, R., Jakobsen, K. S., & Jentoft, S. (2017). De novo gene evolution of antifreeze glycoproteins in codfishes revealed by whole genome sequence data. Mol Biol Evol, 35(3), 593–606.

Baalsrud, H. T., Voje, K. L., Tørresen, O. K., Solbakken, M. H., Matschiner, M., Malmstrøm, M., … Jentoft, S. (2017). Evolution of hemoglobin genes in codfishes influenced by ocean depth. Sci Rep, 7(1), 1–10.

Bankevich, A., Nurk, S., Antipov, D., Gurevich, A. A., Dvorkin, M., Kulikov, A. S., … Prjibelski, A. D. (2012). SPAdes: a new genome assembly algorithm and its applications to single-cell sequencing. Journal of computational biology, 19(5), 455–477. doi:10.1089/cmb.2012.0021

Beck, E. A., Healey, H. M., Small, C. M., Currey, M. C., Desvignes, T., Cresko, W. A., & Postlethwait, J. H. (2021). Advancing human disease research with fish evolutionary mutant models. Trends in Genetics.

Benjamini, Y., & Hochberg, Y. (1995). Controlling the false discovery rate: a practical and powerful approach to multiple testing. Journal of the Royal statistical society: series B (Methodological), 57(1), 289–300.

Betancur-R, R., Broughton, R. E., Wiley, E. O., Carpenter, K., López, J. A., Li, C., … Cureton Ii, J. C. (2013). The tree of life and a new classification of bony fishes. PLoS Curr, 5. doi:10.1371/currents.tol.53ba26640df0ccaee75bb165c8c26288

Bista, I., McCarthy, S. A., Wood, J., Ning, Z., Detrich Iii, H. W., Desvignes, T., … Torrance, J. (2020). The genome sequence of the channel bull blenny, Cottoperca gobio (Günther, 1861). Wellcome Open Research, 5.

Bradnam, K. R., Fass, J. N., Alexandrov, A., Baranay, P., Bechner, M., Birol, I., … Chikhi, R. (2013). Assemblathon 2: evaluating de novo methods of genome assembly in three vertebrate species. GigaScience, 2(1), 2047-2217X-2042-2010.

Brix, O., Clements, K., & Wells, R. (1999). Haemoglobin components and oxygen transport in relation to habitat distribution in triplefin fishes (Tripterygiidae). Journal of Comparative Physiology B, 169(4), 329–334.

Buckner, J. C., Sanders, R. C., Faircloth, B. C., & Chakrabarty, P. (2021). Science Forum: The critical importance of vouchers in genomics. Elife, 10, e68264.

Carrara, N., Weaver, M., Piedade, W. P., Vöcking, O., & Famulski, J. (2019). Temporal characterization of optic fissure basement membrane composition suggests nidogen may be an initial target of remodeling. Developmental biology, 452(1), 43–54.

Chalopin, D., Naville, M., Plard, F., Galiana, D., & Volff, J.-N. (2015). Comparative analysis of transposable elements highlights mobilome diversity and evolution in vertebrates. Genome Biology and Evolution, 7(2), 567–580.

Chawla, H. S., Lee, H., Gabur, I., Vollrath, P., Tamilselvan-Nattar-Amutha, S., Obermeier, C., … Guo, L. (2021). Long-read sequencing reveals widespread intragenic structural variants in a recent allopolyploid crop plant. Plant biotechnology journal, 19(2), 240–250.

Chen, L., DeVries, A. L., & Cheng, C.-H. C. (1997a). Convergent evolution of antifreeze glycoproteins in Antarctic notothenioid fish and Arctic cod. Proceedings of the National Academy of Sciences, 94(8), 3817–3822.

Chen, L., DeVries, A. L., & Cheng, C.-H. C. (1997b). Evolution of antifreeze glycoprotein gene from a trypsinogen gene in Antarctic notothenioid fish. Proceedings of the National Academy of Sciences, 94(8), 3811–3816. doi:10.1073/pnas.94.8.3811

Chen, L., Lu, Y., Li, W., Ren, Y., Yu, M., Jiang, S., … Bilyk, K. T. (2019). The genomic basis for colonizing the freezing Southern Ocean revealed by Antarctic toothfish and Patagonian robalo genomes. GigaScience, 8(4), giz016.

Chénais, B., Caruso, A., Hiard, S., & Casse, N. (2012). The impact of transposable elements on eukaryotic genomes: from genome size increase to genetic adaptation to stressful environments. Gene, 509(1), 7–15.

Cohen, B., Tempelhof, H., Raz, T., Oren, R., Nicenboim, J., Bochner, F., … Ben-Dor, S. (2020). BACH family members regulate angiogenesis and lymphangiogenesis by modulating VEGFC expression. Life science alliance, 3(4).

Cossins, A., & Prosser, C. (1978). Evolutionary adaptation of membranes to temperature. Proceedings of the National Academy of Sciences, 75(4), 2040–2043.

Dainat, J. (2020). AGAT: Another Gff Analysis Toolkit to handle annotations in any GTF/GFF format (Version v0.7.0). Zenodo. doi:10.5281/zenodo.3552717

Davies, P. L., Baardsnes, J., Kuiper, M. J., & Walker, V. K. (2002). Structure and function of antifreeze proteins. Philos Trans R Soc Lond B Biol Sci, 357(1423), 927–935. doi:10.1098/rstb.2002.1081

De Bie, T., Cristianini, N., Demuth, J. P., & Hahn, M. W. (2006). CAFE: a computational tool for the study of gene family evolution. Bioinformatics, 22(10), 1269–1271.

Deng, C., Cheng, C. H., Ye, H., He, X., & Chen, L. (2010). Evolution of an antifreeze protein by neofunctionalization under escape from adaptive conflict. Proc Natl Acad Sci U S A, 107(50), 21593–21598. doi:10.1073/pnas.1007883107

DeVries, A. L., & Steffensen, J. F. (2005). The Arctic and Antarctic polar marine environments. Fish Physiology, 22, 1–24. doi:10.1016/S1546-5098(04)22001-5

di Prisco, G., D’Avino, R., Camardella, L., Caruso, C., Romano, M., & Rutigliano, B. (1990). Structure and function of hemoglobin in Antarctic fishes and evolutionary implications. Polar Biology, 10(4), 269–274.

Ding, F., Li, A., Cong, R., Wang, X., Wang, W., Que, H., … Li, L. (2020). The phenotypic and the genetic response to the extreme high temperature provides new insight into thermal tolerance for the Pacific oyster Crassostrea gigas. Frontiers in Marine Science, 7, 399.

Feng, S., Stiller, J., Deng, Y., Armstrong, J., Fang, Q., Reeve, A. H., … Faircloth, B. C. (2020). Dense sampling of bird diversity increases power of comparative genomics. Nature, 587(7833), 252–257.

Feron, R., Zahm, M., Cabau, C., Klopp, C., Roques, C., Bouchez, O., … Haffray, P. (2020). Characterization of a Y-specific duplication/insertion of the anti-Mullerian hormone type II receptor gene based on a chromosome-scale genome assembly of yellow perch, Perca flavescens. Molecular Ecology Resources, 20(2), 531–543.

Flynn, J. M., Hubley, R., Goubert, C., Rosen, J., Clark, A. G., Feschotte, C., & Smit, A. F. (2020). RepeatModeler2 for automated genomic discovery of transposable element families. Proceedings of the National Academy of Sciences, 117(17), 9451–9457.

Garvin, M. R., Thorgaard, G. H., & Narum, S. R. (2015). Differential expression of genes that control respiration contribute to thermal adaptation in redband trout (Oncorhynchus mykiss gairdneri). Genome Biology and Evolution, 7(6), 1404–1414.

Graham, L. A., Lougheed, S. C., Ewart, K. V., & Davies, P. L. (2008). Lateral transfer of a lectin-like antifreeze protein gene in fishes. PLoS One, 3(7), e2616. doi:10.1371/journal.pone.0002616

Graw, J. (2003). The genetic and molecular basis of congenital eye defects. Nature Reviews Genetics, 4(11), 876–888.

Griffiths, H. J. (2010). Antarctic marine biodiversity–what do we know about the distribution of life in the Southern Ocean? PLoS One, 5(8), e11683.

Hazel, J. R. (1995). Thermal adaptation in biological membranes: is homeoviscous adaptation the explanation? Annual Review of Physiology, 57(1), 19–42.

Heras, J., Chakraborty, M., Emerson, J., & German, D. P. (2020). Genomic and biochemical evidence of dietary adaptation in a marine herbivorous fish. Proceedings of the Royal Society B, 287(1921), 20192327.

Hew, C. L., Wang, N., Joshi, S., Fletcher, G., Scott, G., Hayes, P., … Davies, P. (1988). Multiple genes provide the basis for antifreeze protein diversity and dosage in the ocean pout, Macrozoarces americanus. Journal of Biological Chemistry, 263(24), 12049–12055.

Hobbs, R. S., Hall, J. R., Graham, L. A., Davies, P. L., & Fletcher, G. L. (2020). Antifreeze protein dispersion in eelpouts and related fishes reveals migration and climate alteration within the last 20 Ma. PLoS One, 15(12), e0243273. doi:10.1371/journal.pone.0243273

Hoencamp, C., Dudchenko, O., Elbatsh, A. M., Brahmachari, S., Raaijmakers, J. A., van Schaik, T., … van den Broek, B. (2021). 3D genomics across the tree of life reveals condensin II as a determinant of architecture type. Science, 372(6545), 984–989.

Holborn, M. K., Einfeldt, A. L., Kess, T., Duffy, S. J., Messmer, A. M., Langille, B. L., … Knutsen, T. M. (2021). Reference genome of lumpfish Cyclopterus lumpus Linnaeus provides evidence of male heterogametic sex determination through the AMH pathway. Molecular Ecology Resources.

Holt, C., & Yandell, M. (2011). MAKER2: an annotation pipeline and genome-database management tool for second-generation genome projects. BMC Bioinformatics, 12, 491. doi:10.1186/1471-2105-12-491

Hotaling, S., Borowiec, M. L., Lins, L. S., Desvignes, T., & Kelley, J. L. (2021). The biogeographic history of eelpouts and related fishes: linking phylogeny, environmental change, and patterns of dispersal in a globally distributed fish group. Molecular phylogenetics and evolution, 107211.

Hotaling, S., Kelley, J. L., & Frandsen, P. B. (2021). Towards a genome sequence for every animal: where are we now? Proceedings of the National Academy of Sciences.

Hotaling, S., Sproul, J., Heckenhauer, J., Powell, A., Larracuente, A., Pauls, S., … Frandsen, P. (2021). Long-reads are revolutionizing 20 years of insect genome sequencing. Genome Biology and Evolution, evab138.

Houston, A., & Gingras-Bedard, J. H. (1994). Variable versus constant temperature acclimation regimes: effects on hemoglobin isomorph profile in goldfish, Carassius auratus. Fish physiology and biochemistry, 13(6), 445–450.

Howe, K., Clark, M. D., Torroja, C. F., Torrance, J., Berthelot, C., Muffato, M., … Matthews, L. (2013). The zebrafish reference genome sequence and its relationship to the human genome. Nature, 496(7446), 498–503.

Jones, F. C., Grabherr, M. G., Chan, Y. F., Russell, P., Mauceli, E., Johnson, J., … Kingsley, D. M. (2012). The genomic basis of adaptive evolution in threespine sticklebacks. Nature, 484(7392), 55–61. doi:10.1038/nature10944

Kelley, J. L., Aagaard, J. E., MacCoss, M. J., & Swanson, W. J. (2010). Functional diversification and evolution of antifreeze proteins in the antarctic fish Lycodichthys dearborni. J Mol Evol, 71(2), 111–118. doi:10.1007/s00239-010-9367-6

Kenkel, C. D., & Matz, M. V. (2016). Gene expression plasticity as a mechanism of coral adaptation to a variable environment. Nature Ecology & Evolution, 1(1), 1–6.

Kim, B.-M., Amores, A., Kang, S., Ahn, D.-H., Kim, J.-H., Kim, I.-C., … Lee, J. (2019). Antarctic blackfin icefish genome reveals adaptations to extreme environments. Nature Ecology & Evolution, 3(3), 469–478.

Koren, S., Walenz, B. P., Berlin, K., Miller, J. R., Bergman, N. H., & Phillippy, A. M. (2017). Canu: scalable and accurate long-read assembly via adaptive k-mer weighting and repeat separation. Genome research, 27(5), 722–736.

Korf, I. (2004). Gene finding in novel genomes. BMC Bioinformatics, 5(1), 1–9.

Krueger, F. (2015). Trim Galore!: a wrapper tool around Cutadapt and FastQC to consistently apply quality and adapter trimming to FastQ files. Babraham Bioinformatics, Cambridge, United Kingdom. In.

Lee, S. J., Kim, J.-H., Jo, E., Choi, E., Kim, J., Choi, S.-G., … Park, H. (2021). Chromosomal assembly of the Antarctic toothfish (Dissostichus mawsoni) genome using third-generation DNA sequencing and Hi-C technology. Zoological research, 42(1), 124.

Li, H. (2013). Aligning sequence reads, clone sequences and assembly contigs with BWA-MEM. arXiv, 1303.3997.

Li, H., Handsaker, B., Wysoker, A., Fennell, T., Ruan, J., Homer, N., … Durbin, R. (2009). The sequence alignment/map format and SAMtools. Bioinformatics, 25(16), 2078–2079.

Li, X., Trinh, K.-Y., Hew, C. L., Buettner, B., Baenziger, J., & Davies, P. L. (1985). Structure of an antifreeze polypeptide and its precursor from the ocean pout, Macrozoarces americanus. Journal of Biological Chemistry, 260(24), 12904–12909.

Longo, G. C., Lam, L., Basnett, B., Samhouri, J., Hamilton, S., Andrews, K., … Nichols, K. M. (2020). Strong population differentiation in lingcod (Ophiodon elongatus) is driven by a small portion of the genome. Evol Appl, 13(10), 2536–2554.

Löytynoja, A. (2014). Phylogeny-aware alignment with PRANK. In Multiple sequence alignment methods (pp. 155–170): Springer.

Marks, R. A., Hotaling, S., Frandsen, P. B., & VanBuren, R. (2021). Representation and participation across 20 years of plant genome sequencing. Nature Plants.

Melsted, P., & Pritchard, J. K. (2011). Efficient counting of k-mers in DNA sequences using a bloom filter. BMC Bioinformatics, 12(1), 1–7.

Møller, P. R., Nielsen, J. G., & Anderson, M. E. (2005). Systematics of polar fishes. Fish Physiology, 22, 25–78. doi:10.1016/S1546-5098(04)22002-7

Moore, J. K., Abbott, M. R., & Richman, J. G. (1999). Location and dynamics of the Antarctic Polar Front from satellite sea surface temperature data. Journal of Geophysical Research: Oceans, 104(C2), 3059-3073.

Moran, A. L., & Woods, H. A. (2012). Why might they be giants? Towards an understanding of polar gigantism. Journal of Experimental Biology, 215(12), 1995–2002.

Moran, R. L., Catchen, J. M., & Fuller, R. C. (2020). Genomic resources for darters (Percidae: Etheostominae) provide insight into postzygotic barriers implicated in speciation. Mol Biol Evol, 37(3), 711–729.

Mu, Y., Bian, C., Liu, R., Wang, Y., Shao, G., Li, J., … Ao, J. (2021). Whole genome sequencing of a snailfish from the Yap Trench (~ 7,000 m) clarifies the molecular mechanisms underlying adaptation to the deep sea. PLoS Genet, 17(5), e1009530.

Near, T. J., Dornburg, A., Kuhn, K. L., Eastman, J. T., Pennington, J. N., Patarnello, T., … Jones, C. D. (2012). Ancient climate change, antifreeze, and the evolutionary diversification of Antarctic fishes. Proceedings of the National Academy of Sciences, 109(9), 3434–3439. doi:10.1073/pnas.1115169109

Near, T. J., Parker, S. K., & Detrich III, H. W. (2006). A genomic fossil reveals key steps in hemoglobin loss by the antarctic icefishes. Mol Biol Evol, 23(11), 2008–2016.

Opazo, J. C., Butts, G. T., Nery, M. F., Storz, J. F., & Hoffmann, F. G. (2013). Whole-genome duplication and the functional diversification of teleost fish hemoglobins. Mol Biol Evol, 30(1), 140–153.

Powers, D. A. (1980). Molecular ecology of teleost fish hemoglobins: strategies for adapting to changing environments. American Zoologist, 139–162.

R Core Team. (2021). R: A language and environment for statistical computing.

Rabosky, D. L., Chang, J., Title, P. O., Cowman, P. F., Sallan, L., Friedman, M., … Coll, M. (2018). An inverse latitudinal gradient in speciation rate for marine fishes. Nature, 559(7714), 392–395. doi:10.1038/s41586-018-0273-1

Rice, P., Longden, I., & Bleasby, A. (2000). EMBOSS: the European molecular biology open software suite. Trends in Genetics, 16(6), 276–277.

Robinson, J. T., Thorvaldsdóttir, H., Winckler, W., Guttman, M., Lander, E. S., Getz, G., & Mesirov, J. P. (2011). Integrative genomics viewer. Nature Biotechnology, 29(1), 24–26.

Rothfield, L. I. (2014). Structure and function of biological membranes: Academic Press.

Ruud, J. T. (1954). Vertebrates without erythrocytes and blood pigment. Nature, 173(4410), 848–850.

Sanderson, M. J. (2003). r8s: inferring absolute rates of molecular evolution and divergence times in the absence of a molecular clock. Bioinformatics, 19(2), 301–302.

Scott, G. K., Fletcher, G. L., & Davies, P. L. (1986). Fish Antifreeze Proteins: Recent Gene Evolution. Canadian Journal of Fisheries and Aquatic Sciences, 43(5), 1028–1034. doi:10.1139/f86-128

Scott, G. K., Hew, C. L., & Davies, P. L. (1985). Antifreeze protein genes are tandemly linked and clustered in the genome of the winter flounder. Proc Natl Acad Sci U S A, 82(9), 2613–2617.

Shao, F., Han, M., & Peng, Z. (2019). Evolution and diversity of transposable elements in fish genomes. Sci Rep, 9(1), 1–8.

Shin, S. C., Ahn do, H., Kim, S. J., Pyo, C. W., Lee, H., Kim, M. K., … Park, H. (2014). The genome sequence of the Antarctic bullhead notothen reveals evolutionary adaptations to a cold environment. Genome Biol, 15(9), 468. doi:10.1186/s13059-014-0468-1

Sievers, F., Wilm, A., Dineen, D., Gibson, T. J., Karplus, K., Li, W., … Söding, J. (2011). Fast, scalable generation of high-quality protein multiple sequence alignments using Clustal Omega. Molecular systems biology, 7(1), 539.

Simão, F. A., Waterhouse, R. M., Ioannidis, P., Kriventseva, E. V., & Zdobnov, E. M. (2015). BUSCO: assessing genome assembly and annotation completeness with single-copy orthologs. Bioinformatics, 31(19), 3210–3212.

Smit, A., Hubley, R., & Green, P. (2015). RepeatMasker Open-4.0. 2013–2015. In.

Smit, A. F., & Hubley, R. (2008). RepeatModeler Open-1.0. In.

Sone, J., Mitsuhashi, S., Fujita, A., Mizuguchi, T., Hamanaka, K., Mori, K., … Sugiyama, H. (2019). Long-read sequencing identifies GGC repeat expansions in NOTCH2NLC associated with neuronal intranuclear inclusion disease. Nat Genet, 51(8), 1215–1221.

Stamatakis, A. (2014). RAxML version 8: a tool for phylogenetic analysis and post-analysis of large phylogenies. Bioinformatics, 30(9), 1312–1313. doi:10.1093/bioinformatics/btu033

Stanke, M., & Waack, S. (2003). Gene prediction with a hidden Markov model and a new intron submodel. Bioinformatics, 19(Suppl_2), ii215–ii225.

Star, B., Nederbragt, A. J., Jentoft, S., Grimholt, U., Malmstrom, M., Gregers, T. F., … Jakobsen, K. S. (2011). The genome sequence of Atlantic cod reveals a unique immune system. Nature, 477(7363), 207–210. doi:10.1038/nature10342

Sun, L. F., Chen, X. J., & Jin, Z. B. (2020). Emerging roles of non-coding RNAs in retinal diseases: A review. Clinical & Experimental Ophthalmology, 48(8), 1085–1101.

Supek, F., Bošnjak, M., Škunca, N., & Šmuc, T. (2011). REVIGO summarizes and visualizes long lists of gene ontology terms. PLoS One, 6(7), e21800.

Thomas, G. W., Dohmen, E., Hughes, D. S., Murali, S. C., Poelchau, M., Glastad, K., … Bellair, M. (2020). Gene content evolution in the arthropods. Genome Biology, 21(1), 1–14.

Verde, C., Carratore, V., Riccio, A., Tamburrini, M., Parisi, E., & Di Prisco, G. (2002). The functionally distinct hemoglobins of the Arctic spotted wolffish Anarhichas minor. Journal of Biological Chemistry, 277(39), 36312–36320.

Verde, C., Giordano, D., di Prisco, G., & Andersen, Ø. (2012). The haemoglobins of polar fish: evolutionary and physiological significance of multiplicity in Arctic fish. Biodiversity, 13(3-4), 228-233.

Verde, C., Parisi, E., & di Prisco, G. (2006). The evolution of thermal adaptation in polar fish. Gene, 385, 137–145.

Vona, B., Maroofian, R., Bellacchio, E., Najafi, M., Thompson, K., Alahmad, A., … Shahrokhzadeh, S. (2018). Expanding the clinical phenotype of IARS2-related mitochondrial disease. BMC Med Genet, 19(1), 1–16.

Vurture, G. W., Sedlazeck, F. J., Nattestad, M., Underwood, C. J., Fang, H., Gurtowski, J., & Schatz, M. C. (2017). GenomeScope: fast reference-free genome profiling from short reads. Bioinformatics, 33(14), 2202–2204.

Wang, X., Qu, M., Liu, Y., Schneider, R. F., Song, Y., Chen, Z., … Zhang, S. (2022). Genomic basis of evolutionary adaptation in a warm-blooded fish. The Innovation, 3(1), 100185.

Weber, R. E., Hourdez, S., Knowles, F., & Lallier, F. (2003). Hemoglobin function in deep-sea and hydrothermal-vent endemic fish: Symenchelis parasitica (Anguillidae) and Thermarces cerberus (Zoarcidae). Journal of Experimental Biology, 206(15), 2693–2702.

Wenger, A. M., Peluso, P., Rowell, W. J., Chang, P.-C., Hall, R. J., Concepcion, G. T., … Olson, N. D. (2019). Accurate circular consensus long-read sequencing improves variant detection and assembly of a human genome. Nature Biotechnology, 37(10), 1155–1162.

Willenborg, C., Jing, J., Wu, C., Matern, H., Schaack, J., Burden, J., & Prekeris, R. (2011). Interaction between FIP5 and SNX18 regulates epithelial lumen formation. Journal of Cell Biology, 195(1), 71–86.

Xu, L., Dong, Z., Fang, L., Luo, Y., Wei, Z., Guo, H., … Xia, Q. (2019). OrthoVenn2: a web server for whole-genome comparison and annotation of orthologous clusters across multiple species. Nucleic Acids Res, 47(W1), W52-W58.

Xu, S., Wang, J., Guo, Z., He, Z., & Shi, S. (2020). Genomic convergence in the adaptation to extreme environments. Plant communications, 100117.

Yang, Z. (2007). PAML 4: phylogenetic analysis by maximum likelihood. Mol Biol Evol, 24(8), 1586–1591.

Zhuang, X., Yang, C., Murphy, K. R., & Cheng, C.-H. C. (2019). Molecular mechanism and history of non-sense to sense evolution of antifreeze glycoprotein gene in northern gadids. Proceedings of the National Academy of Sciences, 116(10), 4400–4405.

