## Supporting Information for "Pathways to polar adaptation in fishes revealed by long-read sequencing"

**Contents:**

Figures S1 – S4

Table S1

Tables S2-S5 provided as an Excel table

**Figures:**

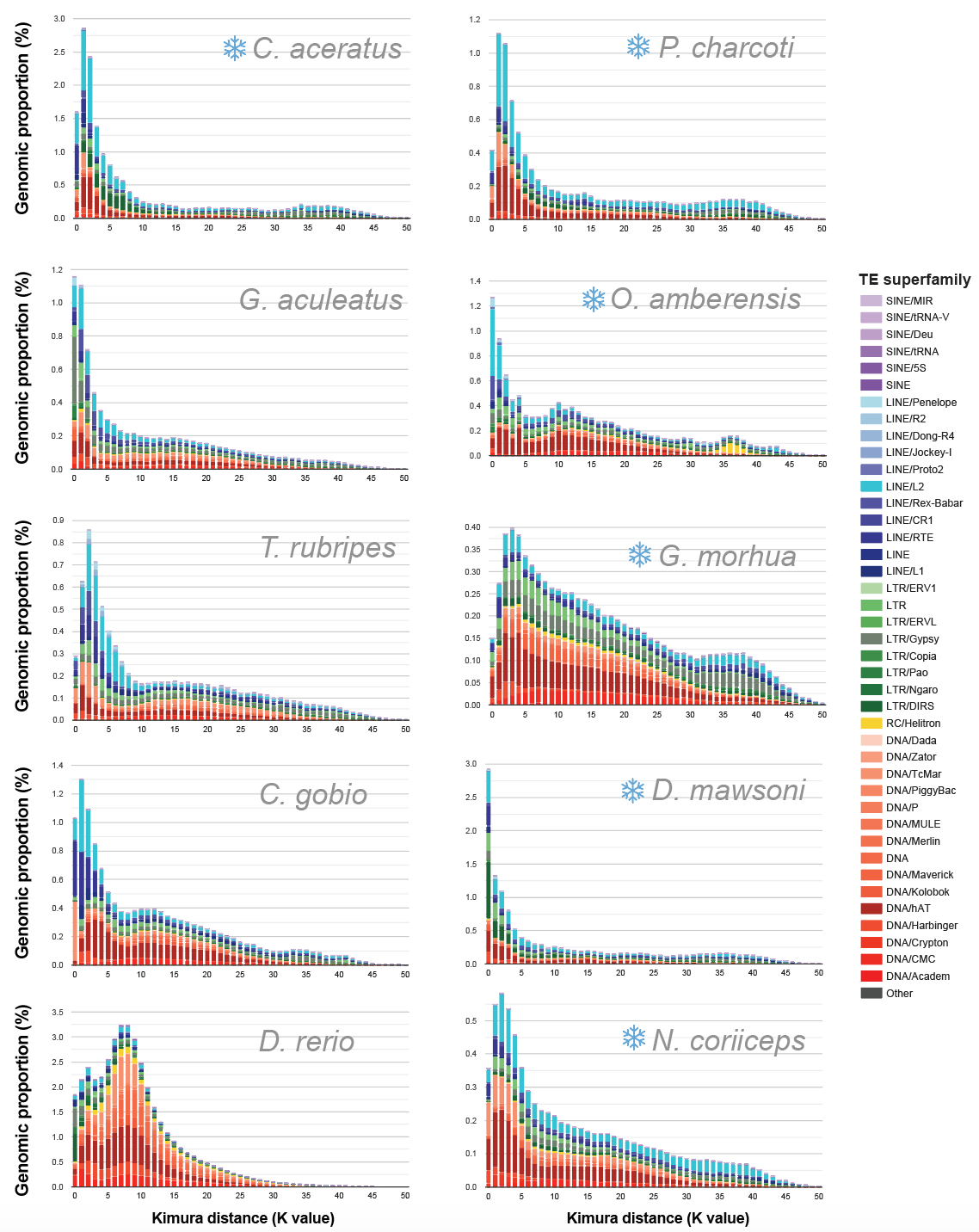

**Figure S1.** **TE superfamily copy divergence**. Transposable element (TE) superfamily across the focal species included in this study. Snowflakes indicate species with polar distributions. The y-axis shows TE abundance as a proportion of the genome (e.g., 1.0 = 1% of the genome). The x-axis shows sequence divergence (CpG adjusted Kimura distance) relative to consensus sequences for TE superfamilies. Copy number peaks with abundance skewed toward the left (i.e., low sequence divergence) may represent TE copies with a recent history of diversification relative to peaks with right-skewed abundance which may comprise remnants of more ancient bursts of TE activity. TE superfamilies are separated by color to show additional classification details in support of Fig. 2.

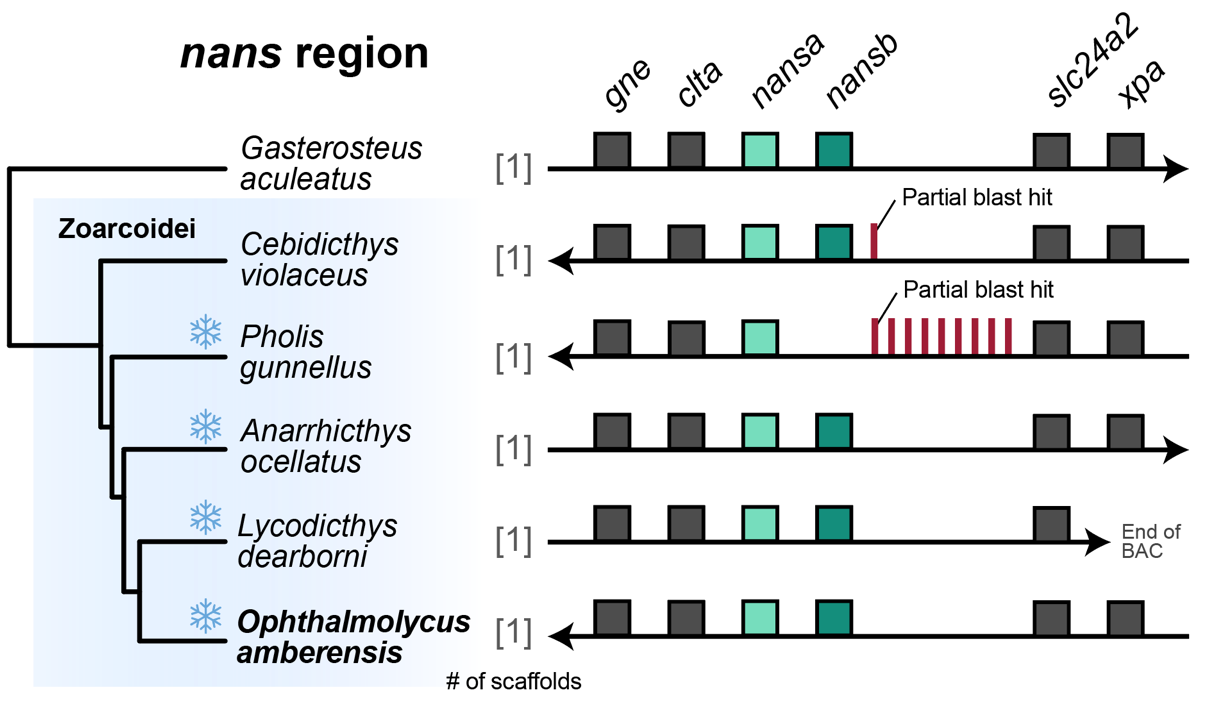

**Figure S2.** Synteny of the sialic acid synthase (*nans*) region in the suborder Zoarcoidei with three-spined stickleback (*G. aculeatus*) as the outgroup. Snowflakes indicate species with distributions that overlap polar regions. The arrowheads indicate the direction of the clusters in each assembly. The position of each square above or below the line denotes the position of the gene on the forward (when the arrow points to the right) or reverse strand (when the arrow points to the left), respectively.

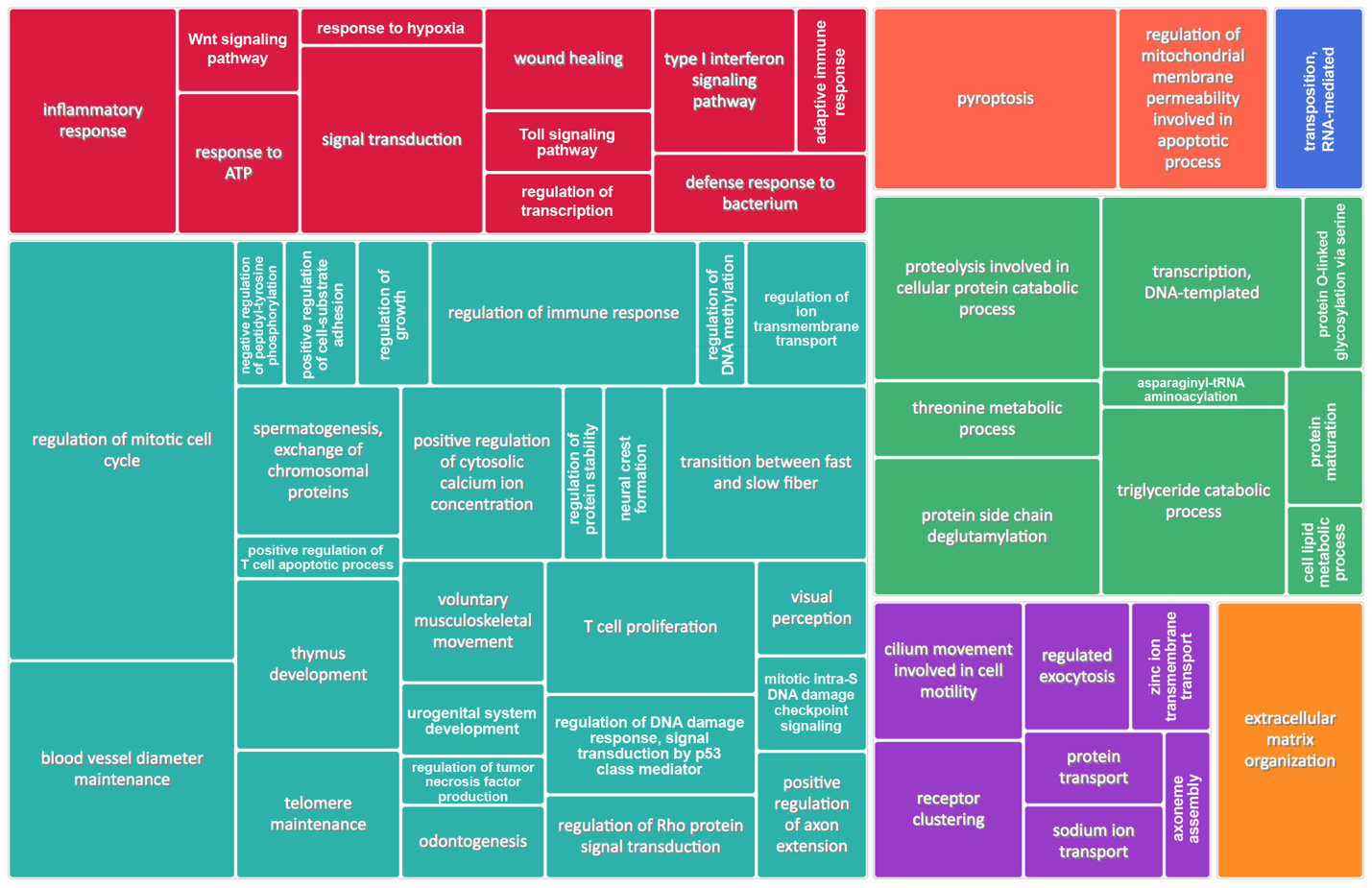

**Figure S3.** Gene ontology terms of rapidly expanding gene families in *O. amberensis*. Colors reflect general functional categories and box sizes reflect the scale of gene family expansion for each GO term.

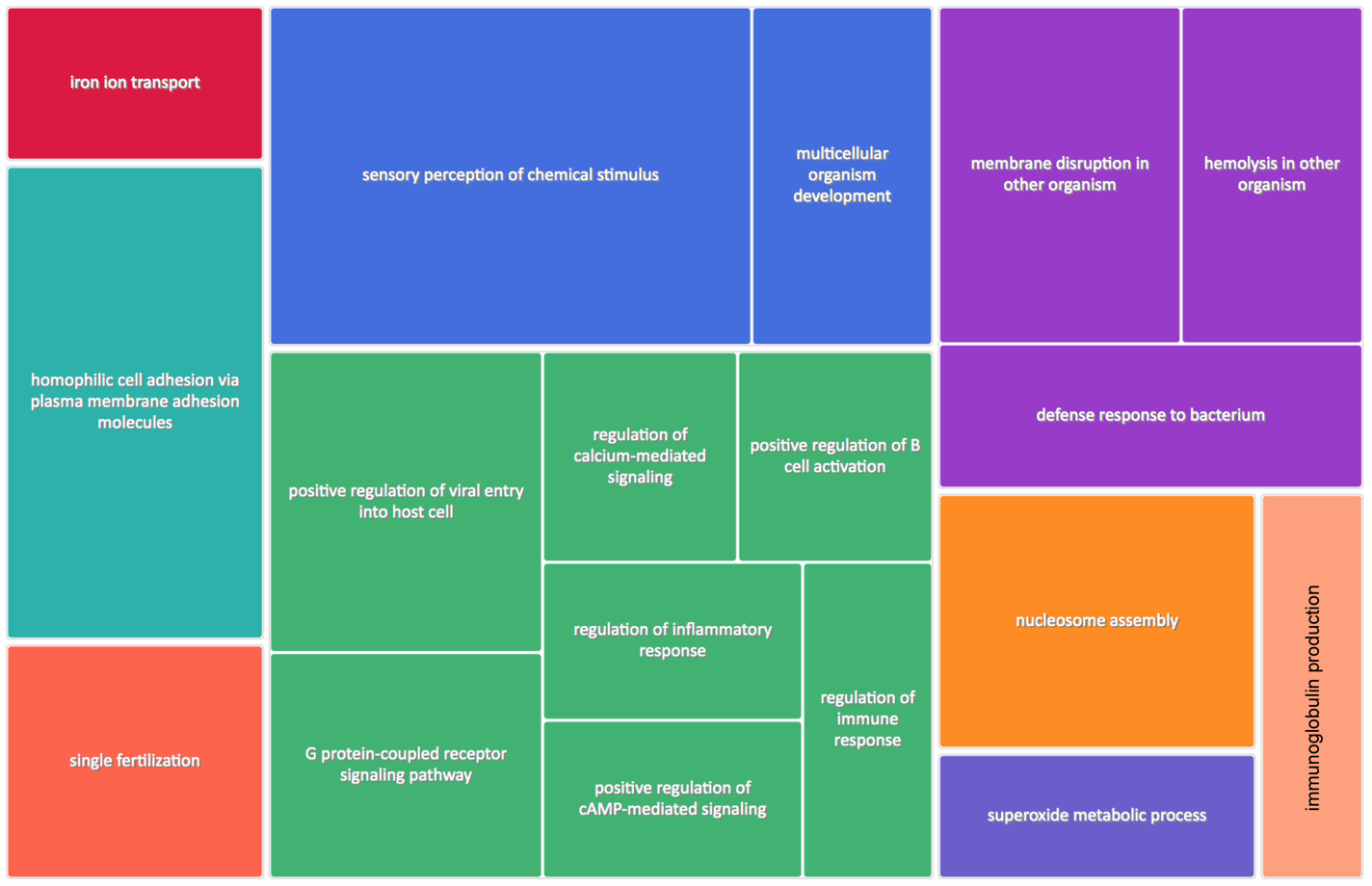

**Figure S4.** Gene ontology terms of rapidly contracting gene families in *O. amberensis*. Colors reflect general functional categories and box sizes reflect the scale of gene family contraction for each GO term.

**Tables:**

**Table S1.** Genome assembly metrics for the species that were included in either the hemoglobin synteny analyses (Figure 4) or *afpIII* synteny analyses (Figure 5). *Lycodicthys dearborni* is not included because only BACs were used.

| Species | Assembled Genome Length | Number of scaffolds | Longest scaffold | Scaffold N50 | %N, assembly |
| --- | --- | --- | --- | --- | --- |
| *P. flavescens* | 877.46 Mb | 268 | 44.58 Mb | 37.41 Mb | 0.05% |
| *E. spectable* | 854.79 Mb | 3,118 | 38.81 Mb | 30.50 Mb | 0.47% |
| *C. gobio* | 609.39 Mb | 322 | 30.48 Mb | 25.16 Mb | 0.42% |
| *G. aculeatus* | 461.53 Mb | 1,844 | 32.63 Mb | 18.11 Mb | 3.23% |
| *C. violaceus* | 593.00 Mb | 467 | 19.93 Mb | 6.72 Mb | 0.01% |
| *P. gunnellus* | 588.73 Mb | 102 | 31.68 Mb | 25.36 Mb | 0.02% |
| *A. ocellatus* | 612.79 Mb | 10,817 | 19.78 Mb | 5.72 Mb | 7.15% |
| *O. amberensis* | 679.26 Mb | 1,828 | 6.97 Mb | 1.00 Mb | 0% |
| *O. elongatus* | 645.89 Mb | 2,175 | 13.37 Mb | 1.75 Mb | 0% |
| *C. lumpus* | 572.90 Mb | 49 | 31.50 Mb | 23.86 Mb | 1.78% |
| *Pseudoliparis sp.* Yap Trench | 840.64 Mb | 3,134 | 8.36 Mb | 0.98 Mb | 0.50% |
| *T. bubalis* | 615.15 Mb | 27 | 50.13 Mb | 29.14 Mb | 0.01% |

**Table S2.** See additional Excel file.

**Table S3.** See additional Excel file.

**Table S4.** See additional Excel file.

**Table S5.** See additional Excel file.
